# Dissecting the transcriptomic basis of phenotypic evolution in an aquatic keystone grazer

**DOI:** 10.1101/606988

**Authors:** Dagmar Frisch, Dörthe Becker, Marcin W. Wojewodzic

## Abstract

Cultural eutrophication is one of the largest threats to aquatic ecosystems with devastating ecological and economic consequences. Although *Daphnia*, a keystone grazer crustacean, can mitigate some negative effects of eutrophication in freshwater habitats, it is itself affected by changes in nutrient composition. Previous studies have shown evolutionary adaptation of *Daphnia* to environmental change, including high phosphorus levels. While phosphorus is an essential macronutrient, it negatively affects *Daphnia* life history when present in excessive concentrations. Here, we map weighted gene co-expression networks to phosphorus-related phenotypic traits in centuries-old ancestral *Daphnia* and their modern descendants, contrasting pre- and post-eutrophication environments. We find that evolutionary fine-tuning of transcriptional responses is manifested at a basic (cellular) phenotypic level, and is strongly correlated to an evolved plasticity of gene expression. At a higher phenotypic level, this contributes to the maintenance of physiological homeostasis, and a robust preservation of somatic phosphorus concentration.

## Introduction

Cultural eutrophication is one of the largest threats to aquatic ecosystems with devastating ecological and economic effects^1^. Rising phosphorus (P) concentrations in surface water result in high algal biomasses that pose a serious ecological threat to aquatic habitats, impacting biodiversity, drinking water quality, recreational resources, and fisheries^1-2^. One of the major P-sources of eutrophication is the use of phosphate fertilizers, for which the global consumption has increased from 1 to 15 Mt P year^-1^ in the 20th century^3^. Vital ecosystem services provided by *Daphnia*, a keystone herbivore crustacean in freshwater habitats, can effectively mitigate some of these negative impacts by decimating phytoplankton biomass^4^. Although P is a vital element for many cellular components, including RNA, DNA and ATP, excessive P-concentrations typical of eutrophication can negatively affect somatic and population growth of *Daphnia*^5-6^. In addition, algal blooms often involve toxin-producing cyanobacteria that can negatively affect the growth rate of *Daphnia*, although local adaptation to these toxins has been observed^7-9^. Insights from resurrection ecology^10^ suggest that *Daphnia* can adapt to changing environmental conditions, such as climate change^11^, anthropogenic stressors^12^, toxic algae^13^, and phosphorus supply^14^. Although previous analyses have examined the functional genome underlying *Daphnia* phenotypes (e.g.^15-19^), the link between phenotypic adaptation and the evolution of gene expression reaction norms in this keystone species is poorly understood.

Adaptation to environmental change has been associated with divergent gene expression patterns^20^ and the maintenance or evolution of gene expression reaction norms^21-22^. However, directly linking phenotypic trait evolution and its molecular drivers can be daunting because of the complex transcriptional architecture of gene expression and associated regulatory processes^23^. In order to account for such complexity, novel approaches and tools can now be used to elucidate the genetic basis of phenotypic traits^24-27^. Especially, co-expression gene network analyses have provided key insights to the molecular basis of phenotypic evolution^28^, and adaptive phenotypic plasticity^29^.

Here, using ‘resurrection ecology’, we evaluate transcriptomic responses to P availability of ancient and modern *Daphnia* associated with three phosphorus-related phenotypic traits (P retention rate, somatic P content, and somatic growth rate^14,30^). We do so by mapping weighted gene co-expression networks to traits previously assessed in resurrected ancient (600 years-old) and modern *Daphnia* from a lake with a 24-fold historic change in sediment P-flux^14,31^. In a second step we assess evolutionary conservation or divergence in transcriptional networks^32-33^ of the same isolates. Together, the employed analyses allowed us, for the first time in an ecological keystone species, to decipher genetic key drivers of phenotypic evolution. Our study highlights that phenotypic evolution is a result of molecular fine-tuning on different layers ranging from basic cellular responses to higher order phenotypes. In the context of eutrophication, these findings advance our understanding how populations are able to persist throughout major environmental shifts and to maintain their vital ecosystem services.

## Results

### Trait-associated network

Separation of gene expression profiles of ancient (ancestral) and modern *Daphnia* by evolutionary history and phosphorus (P)-availability (high or low P supply) (Fig. 1a) suggests divergent transcriptional regulation between ancestral clones and their descendants. Module membership of all 10,439 genes that entered the analysis was resolved, and resulted in 17 colour-coded (Fig. 1b, details in Table S1, Fig. S1) and clearly defined modules (e.g., Fig. 1c). The obtained gene co-expression network was used to quantitatively link gene expression to three phenotypic traits associated with P-physiology: P-retention efficiency (RE), somatic (body) P content (bP), and somatic growth rate (GR). Previous studies showed that phenotypic responses of RE and GR differed significantly between ancient and modern isolates^14^ while their bP was similar^30^. Those studies suggested the evolution of phenotypic plasticity related to nutrient enrichment, in particular of RE^14^ (Fig. 2a). RE did not directly translate to bP (Fig. 2a), indicating a decoupling of these two traits as observed in other studies^30,34-35^. GR of ancient *Daphnia* was similar in both P-treatments, while GR of modern *Daphnia* was strongly determined by P-availability, suggesting a clear association with physiological traits such as RE and bP^36^.

**Fig 1.**
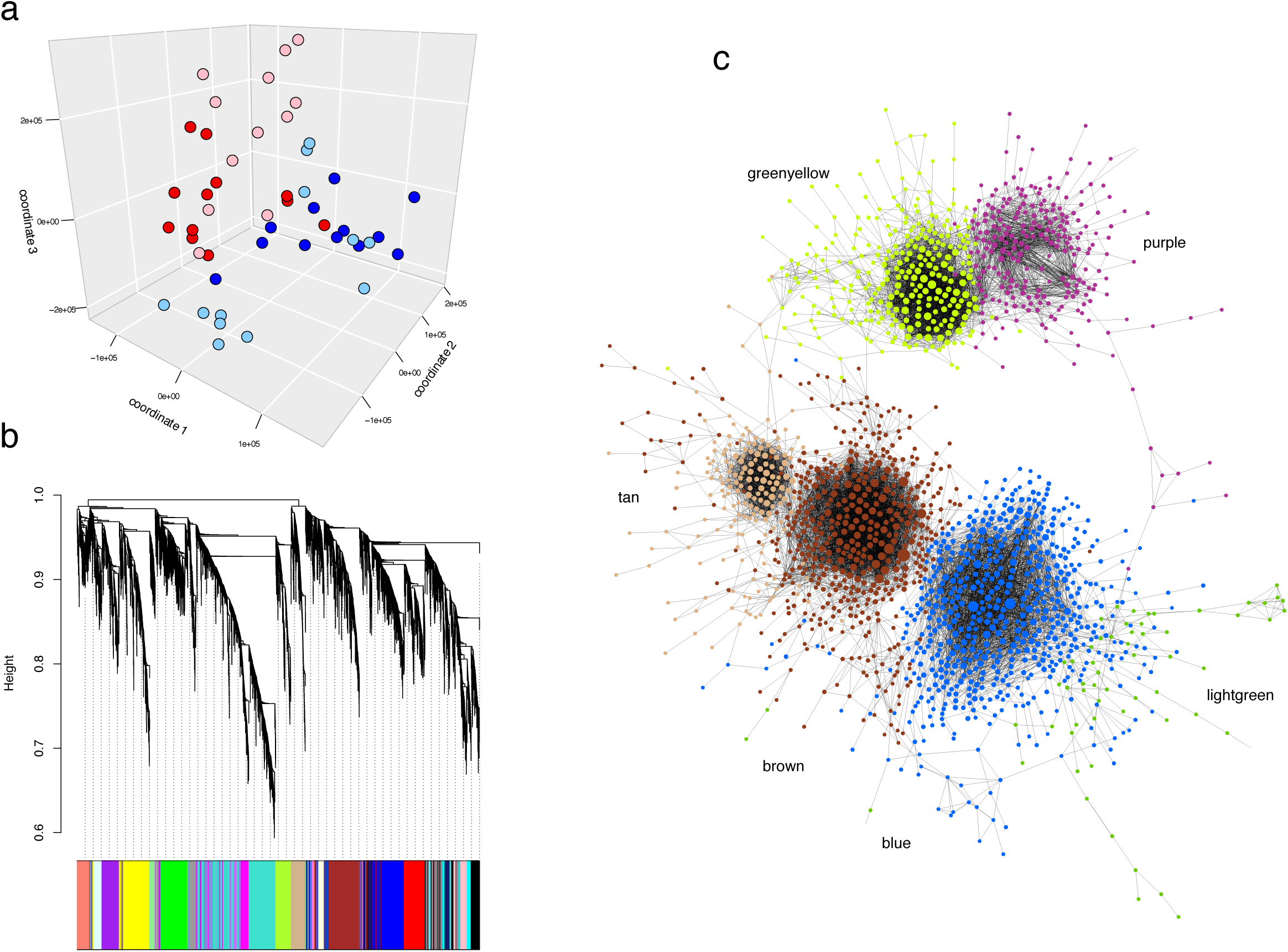
Visualization of gene expression profiles and the trait-associated gene co-expression network of four resurrected *Daphnia pulicaria* clones. a. Multi dimensional scaling (MDS) plot of average gene expression profiles of four resurrected *Daphnia pulicaria* clones in response to P-supply. Red symbols: ancient clones, blue symbols: modern clones, lighter shading: low P food, darker shading: high P food. b. Gene dendrogram resulting from average linkage hierarchical clustering, and identified modules (colour bar). c. Gene network of six modules correlated with one of the measured phenotypic traits. Nodes represent individual genes present in the modules, with their size relative to within-module connectivity (larger nodes have a higher connectivity). To reduce complexity of the network representation, only nodes are shown that are connected by edges (intramodular connectivity k) with a weight >0.4.

**Fig. 2.**
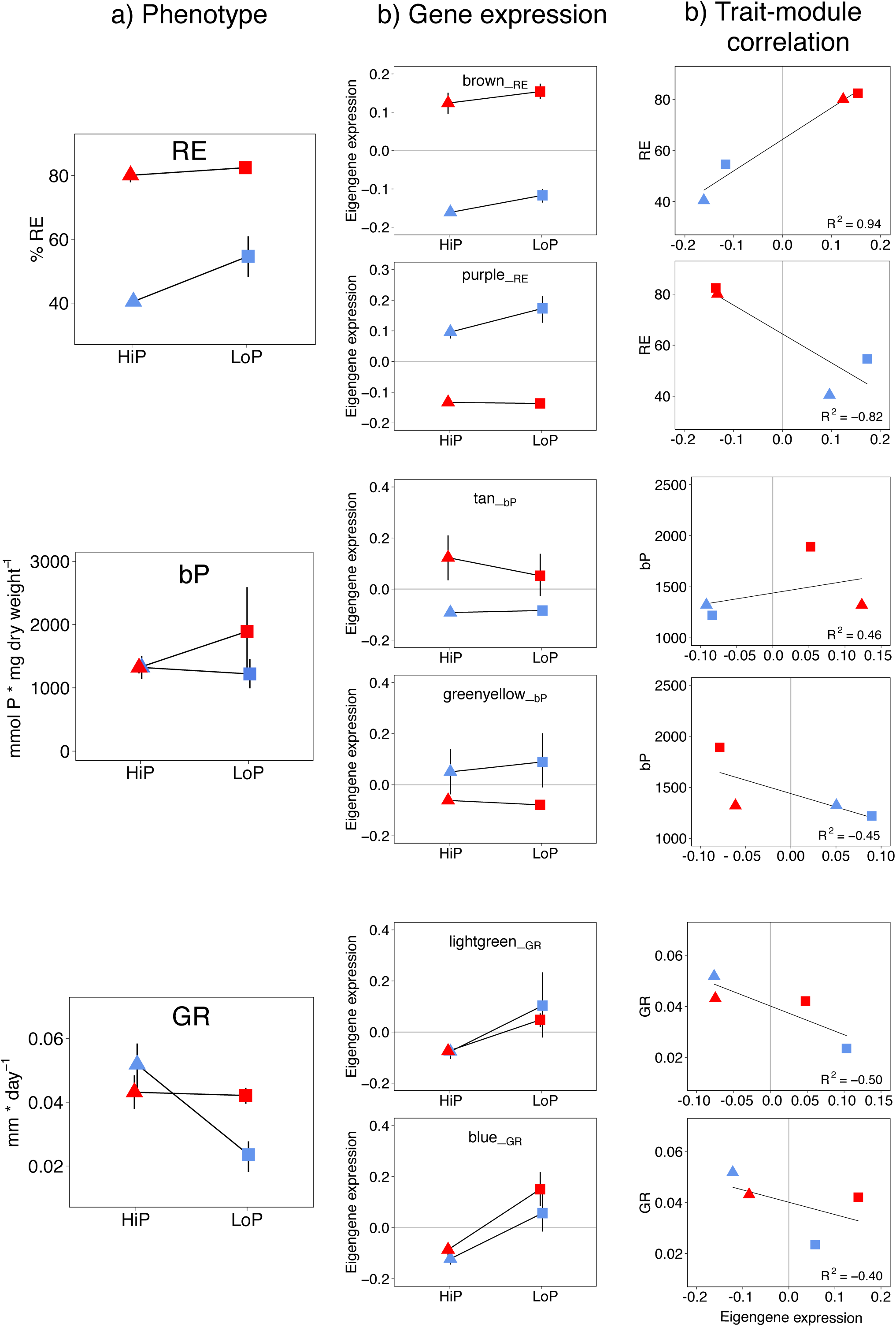
Relationship between phenotypic traits and correlated modules identified in the trait-associated network. Error bars represent 95% confidence intervals. a. Phenotypic trait expression at high (HiP; triangles) and low (LoP; squares) phosphorus supply. RE = retention efficiency, GR = growth rate, bP = body P content. b. Eigengene expression of the two modules most highly positively or negatively correlated with one of the measured phenotypic traits. c. Correlation between phenotypic trait (y axis) and eigengene expression (x-axis); R^2^ values are computed values of the WGCNA network statistics.

### Module-trait correlation and functional analysis

Our analysis identified groups of genes (i.e., modules) as drivers of the observed phenotypic variation. Here, we only consider in detail the two most highly correlated modules per trait (Fig. 1c, Fig S2). These six modules together contained about one third of the input gene set. *Note*: For readability, the six modules correlated with one of the three phenotypic traits will in the following be referred to as trait-modules and denoted as *module-colour*__ TRAIT_ (e.g., *blue*__GR_).

### Retention efficiency (RE)

The phenotypic response of ancient clones to P-supply suggests canalisation of the RE trait, while that of modern clones suggests the evolution of phenotypic plasticity^14^ (Fig. 2a). These phenotypic trait values were reflected in the magnitude of gene expression of the two modules most highly correlated with RE: the positively correlated *brown*__RE_ and the negatively correlated *purple*__RE_ module (Fig. 2b, c). Overall, *brown*__RE_ genes were more highly expressed in ancient clones, whereas *purple*__RE_ genes were more highly expressed in modern clones. This pattern suggests involvement of *brown*__RE_ genes in the overall higher RE of ancient clones.

Functionally, *brown*__RE_ was enriched in cellular processes and signalling (Fig. 3a). Other modules positively correlated with RE (Fig. S2-S4) were additionally enriched in metabolic processes, mainly involving lipid transport. The functional enrichment of *brown*__RE_ was reflected in genes controlled by the main promoter motifs of this module _with_ many of them involved in cellular processes (Fig. 3c). Two of the three most strongly connected genes (i.e., hubgenes) of *brown*__RE_ belonged to the *jumonji* gene family that is known to regulate chromatin organization and thus gene expression^37^. This finding suggests epigenetic regulation, previously described as DNA compaction in *Daphnia*^38^. The third hubgene as well as 50% of the *brown*__RE_ genes in general lacked functional annotation (Fig. 3b, Fig. S5a).

**Fig. 3.**
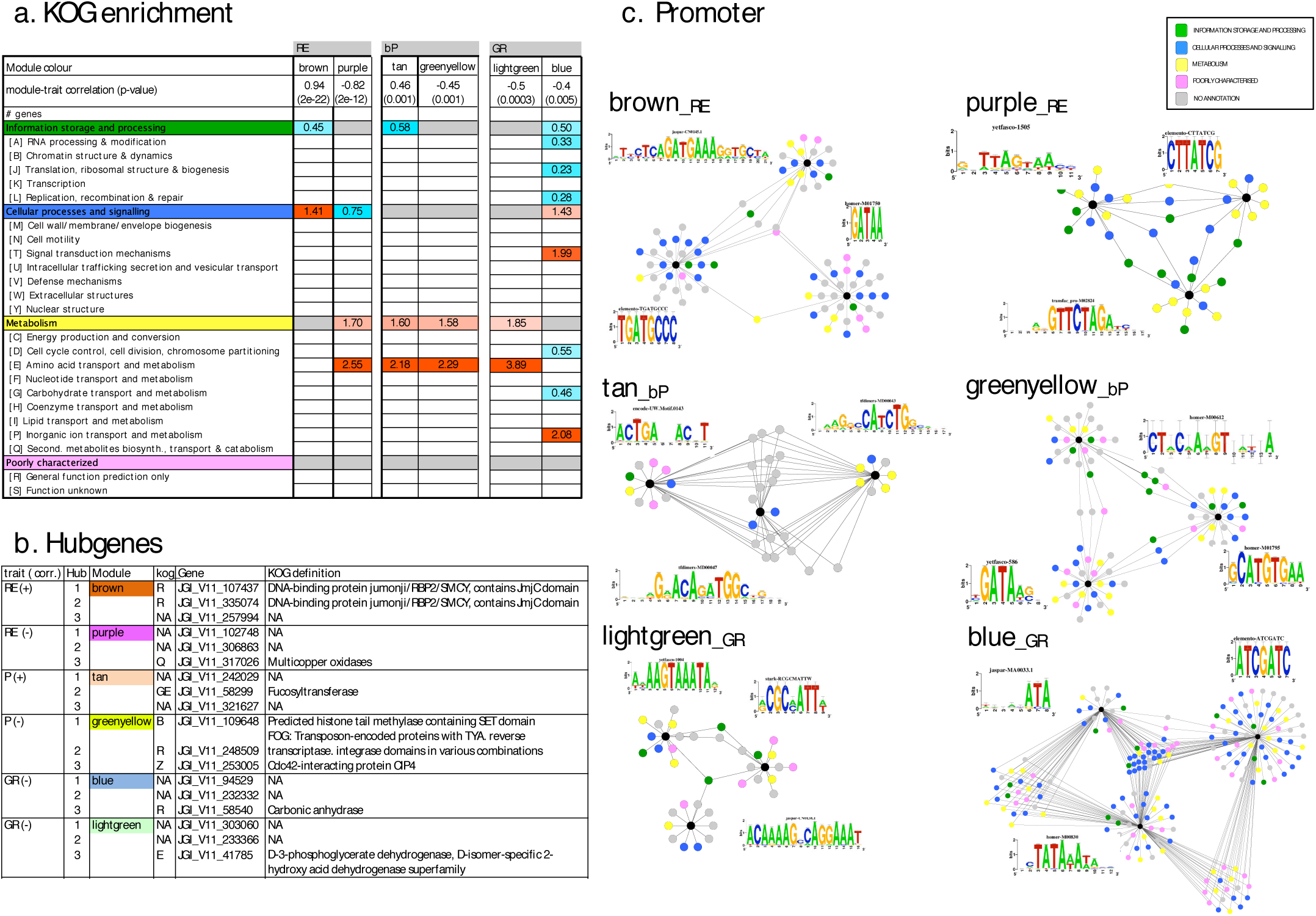
Functional analysis of the six modules most highly correlated (R^2^ > 0.4, p ≤ 0.005) with one of the phenotypic traits. a. Functional enrichment (using KOG categories) of modules. Enrichment scores (odds-ratios, FDR <0.05) are calculated separately for each module and can therefore only be directly compared within columns. b. Top three hubgenes of the trait-associated modules. c. Cytoscape networks of potential promoter target genes. Colour coding (i.e., green, blue, yellow, pink and grey) indicates functional association of nodes to distinct KOG categories (for details see insert). Black nodes represent promoter motifs binding sites (also shown as sequence logo).

In modern clones, a strong positive association between *purple*__RE_ gene expression and RE (Fig. 2a,b) indicates an important role of these genes in trait plasticity. *Purple*__RE_ was functionally enriched in metabolic processes, specifically in amino acid transport and metabolism (Fig. 3a) including several genes coding for trypsin and chitinase that were more highly expressed in modern clones, especially in low P conditions (Fig. S5b). *Purple*__RE_ genes controlled by the main promoter motifs also included several trypsins and other genes involved in metabolic processes (Fig. 3c). Overrepresentation of genes involved in aminoacid metabolism (including trypsins) may indicate the exploitation of alternate P-sources under P-limitation as seen in plants^39^. Other modules negatively correlated with RE (Fig. S4) were additionally enriched in cell cycle control, cell division and carbohydrate transport as well as processes involved in information storage and processing.

One of the three top hubgenes of *purple*__RE_ codes for a multi-copper oxidase (MCO) (Fig. 3b), a gene family essential for iron metabolism in many organisms^40^. Previous research in *Daphnia* identified a significant interaction between P-limitation and iron-kinetics^41^. This finding, together with our results suggests a central role of MCOs in modulating essential cellular processes under P-limitation. Both other top hubgenes as well as about 30% of *purple*__RE_ genes lacked functional annotation (Fig. S5b).

### Body P (bP)

Body P was significantly correlated with eigengene expression of *tan*__bP_ and *greenyellow*__bP_ (Fig. 2b,c). Genes in *tan*__bP_ were overall more highly expressed in ancient clones, with higher values in the high P (HiP) treatment. Generally, *tan*__bP_ genes were more highly expressed in ancient clones, and enriched in amino acid metabolism (Fig. 3a, Fig. S5c), with one of the hubgenes identified as a fucosyltransferase, an enzyme involved in glycosylation (Fig. 3b).

We speculate that the *tan*__bP_ genes highly expressed under HiP in ancient clones including many non-annotated genes (Fig. S5c) may contribute to the maintenance of defined bP concentrations under these conditions. Such a regulation might be necessary in order to counteract the effect of unusually high RE in ancient clones, and to retain cellular homeostasis, e.g. by active release of inorganic phosphorus^42^or moulting^43^. When evaluating the genes controlled by the top three promoter motifs, we found many non-annotated genes of which the majority seems to be controlled simultaneously by two or even three promoter motifs, indicating a high degree of co-regulation (Fig. 3c).

Genes in *greenyellow*__bP_ were overall more highly expressed in modern clones, with higher values in the low P (LoP) treatment. Similar to *tan*__bP_, *greenyellow*__bP_ was enriched for processes involved in amino acid transport and metabolism (Fig. 3a). This functional pattern was reflected in the genes controlled by the top promoter motifs with many of them involved in metabolic processes (Fig. 3c); about 40% of *greenyellow*__bP_ genes had no functional annotation (Fig. S5d). Identified hubgenes (Fig. 3c) included a predicted histone tail methylase (an epigenetic modifier), a FOG gene (putative cell signal^44^) and a Cdc42 (a conserved protein involved in cell cycle regulation^45^). Histone tail methylation has profound effects on gene transcription and can be passed transgenerationally in invertebrates, with the possibility of a long-lasting epigenetic memory of environmental conditions^46^. Correlation of bP with genes involved in protein metabolism suggest that these genotypes are able to maintain homeostasis in body P-content in response to dietary P-supply by producing metabolic adjustments in P-usage.

### Somatic growth rate (GR)

According to the Growth Rate Hypothesis^36,47^, GR is a trait that strongly depends on various molecular and physiological parameters controlling P-allocation to ribosomal RNA. The two highest trait-module correlations of GR were detected with *lightgreen*__GR_ and *blue*__GR_ (Fig. 2c). Functionally, *lightgreen*__GR_ was enriched for metabolic processes, specifically amino acid transport and metabolism (Fig. 3a) including several trypsin-coding genes and several genes related to P-metabolism. *Blue*__GR_ was enriched in signal transduction mechanisms including a number of genes encoding phosphodiesterases and phosphatases. Genes of this functional gene family are known to be involved in P-scavenging in plants^48^, but also in *Daphnia* (*e.g.,* alkaline phosphatase^49^). About 30% and 50% of the GR-associated genes (*blue*__GR_ and *lightgreen*__GR_, respectively) lacked functional annotation (Fig. S5e,f). Gene expression in both modules was negatively correlated with GR, and enriched in amino acid and inorganic ion transport and metabolism that could increase P-availability and uptake under P-starvation. However, GR was also correlated with two other modules, albeit more weakly, that were enriched in functional groups more directly related to somatic growth such as transcription, translation, and replication and cell cycle control (turquoise and magenta modules, Figs. S2-S4). In contrast to the trait-module correlations of RE and bP, which were driven by evolutionary history (ancient or modern clones), GR module correlations were determined by treatment, with similar responses of ancestral and derived clones, suggesting environmentally induced gene expression. The observed functional enrichment in signalling cascades involving transmitters and receptors indicates such environmental triggering of gene expression, particularly in *blue*__GR_. When P supply in the environment is not limiting, these signalling cascades may lead to an increased rRNA biogenesis thus increasing GR^36^. Co-regulation of these genes by several transcription factors lends further support to this idea: genes controlled by the top three promoter motifs in each of the two modules reflect the same functional enrichment as the entire module (Fig. 3c). Notably, a large number of *blue*__GR_ genes are potentially co-regulated by two or more promoter binding sites (Fig. 3c). In both GR-modules only one hubgene was functionally annotated, namely genes coding for the enzymes phosphoglycerate dehydrogenase (PHGDH) in *lightgreen*__GR_ and carbonate dehydratase (carbonic anhydrase, CA) in *blue*__GR_ (Fig. 3c). PHGDH is crucial to L-serine biosynthesis, an amino acid central to cellular proliferation^50^, and CA is a zinc metalloenzyme involved in several biological processes such as the transport of CO_2_, maintaining acid-base balance, glycogen and lipid synthesis^51^. Both enzymes therefore mediate multiple pathways that can play a significant role in regulating growth.

#### Gene expression plasticity

All six trait-modules in focus showed some degree of gene expression plasticity in either ancestral or derived clones (Fig. 2b). In one of the two trait-modules associated with RE (*purple*__RE_), gene expression of the derived clones was significantly plastic but not in the ancestral clones (Table 1). A similar pattern was observed for *greenyellow*__bP_. The other module associated with bP (*tan*__bP_) showed significant plasticity in response to P-supply in the ancient clones but not in the derived *Daphnia*. Gene expression of both ancestral and derived *Daphnia* was significantly plastic in response to P-supply in the two GR-related modules (*blue*__GR_ and *lightgreen*__GR_) and in one of the modules associated with RE (*brown*__RE_).

**Table 1.** Results of Linear Mixed Effect models for Module Eigengene (ME) expression with evolutionary history (‘era’: ancient or modern) and food treatment (high P (HiP) or low P (LoP) algae) as fixed effects, and clone (two ancient and two modern clones) as random factor, nested in ‘era’.

| Module |  | Estimate | SE | df | t value | Pr(>\|t\|) |  |
| --- | --- | --- | --- | --- | --- | --- | --- |
| *brown*_RE | (Intercept) | 0.891 | 0.009 | 48 | 95.251 | 2.34E-56 | *** |
|  | treatLP | -0.024 | 0.013 | 48 | -1.819 | 0.075 |  |
|  | eraM | 0.301 | 0.013 | 48 | 22.772 | 2.24E-27 | *** |
|  | treatLP:eraM | -0.035 | 0.019 | 48 | -1.879 | 0.066 |  |
| *purple*_RE | (Intercept) | 1.778 | 0.037 | 48 | 47.900 | 3.48E-42 | *** |
|  | treatLP | 0.027 | 0.052 | 48 | 0.516 | 0.608 |  |
|  | eraM | -1.077 | 0.052 | 48 | -20.520 | 2.10E-25 | *** |
|  | treatLP:eraM | -0.178 | 0.074 | 48 | -2.395 | 0.021 | * |
| *tan*_bP | (Intercept) | 0.909 | 0.067 | 4.0 | 13.525 | 1.62E-04 | *** |
|  | treatLP | 0.063 | 0.010 | 44.0 | 6.393 | 8.92E-08 | *** |
|  | eraM | 0.193 | 0.095 | 4.0 | 2.030 | 0.111 |  |
|  | treatLP:eraM | -0.072 | 0.014 | 44.0 | -5.179 | 5.33E-06 | *** |
| greenyellow_bP | (Intercept) | 1.297 | 0.276 | 4.0 | 4.702 | 0.009 | ** |
|  | treatLP | 0.093 | 0.037 | 44.0 | 2.516 | 0.016 | * |
|  | eraM | -0.257 | 0.390 | 4.0 | -0.658 | 0.546 |  |
|  | treatLP:eraM | -0.208 | 0.052 | 44.0 | -3.990 | 2.47E-04 | *** |
| *lightgreen*_GR | (Intercept) | 1.169 | 0.116 | 4.4 | 10.040 | 3.44E-04 | *** |
|  | treatLP | -0.252 | 0.049 | 44.0 | -5.135 | 6.18E-06 | *** |
|  | eraM | 0.016 | 0.165 | 4.4 | 0.099 | 0.926 |  |
|  | treatLP:eraM | -0.002 | 0.069 | 44.0 | -0.029 | 0.977 |  |
| *blue*_GR | (Intercept) | 0.956 | 0.018 | 6.3 | 51.857 | 1.54E-09 | *** |
|  | treatLP | 0.116 | 0.017 | 44.0 | 6.913 | 1.53E-08 | *** |
|  | eraM | -0.019 | 0.026 | 6.3 | -0.716 | 0.500 |  |
|  | treatLP:eraM | -0.026 | 0.024 | 44.0 | -1.119 | 0.269 |  |

#### Module preservation in ancient and modern gene co-expression networks

Construction of gene co-expression networks separately for ancient and modern *Daphnia* (Fig. S6) allowed the computation of network statistics that revealed moderate to strong preservation of ∼ 70% of modules detected in ancient clones (Fig. S7). *Note*: a module is considered *preserved* if their network structures are similar (see methods for details), yet their corresponding gene sets may not strongly overlap^52^. To avoid ambiguity, we distinguish modules of the preservation networks from those of the trait-associated network by denoting them as *module-colour*_Pres_ (e.g. *yellow*_Pres_). Our results generally support strong evolutionary conservation of gene expression patterns in ancient and modern *Daphnia* clones despite their temporal separation over centuries. However, evolutionary modification of transcriptional patterns was indicated in a small number of moderately to non-preserved modules (Fig. S7). In contrast to non-preserved modules, functional enrichment in each of the most highly preserved modules was similar in ancient and modern clones (Fig. 5 and S8).

**Fig. 4.**
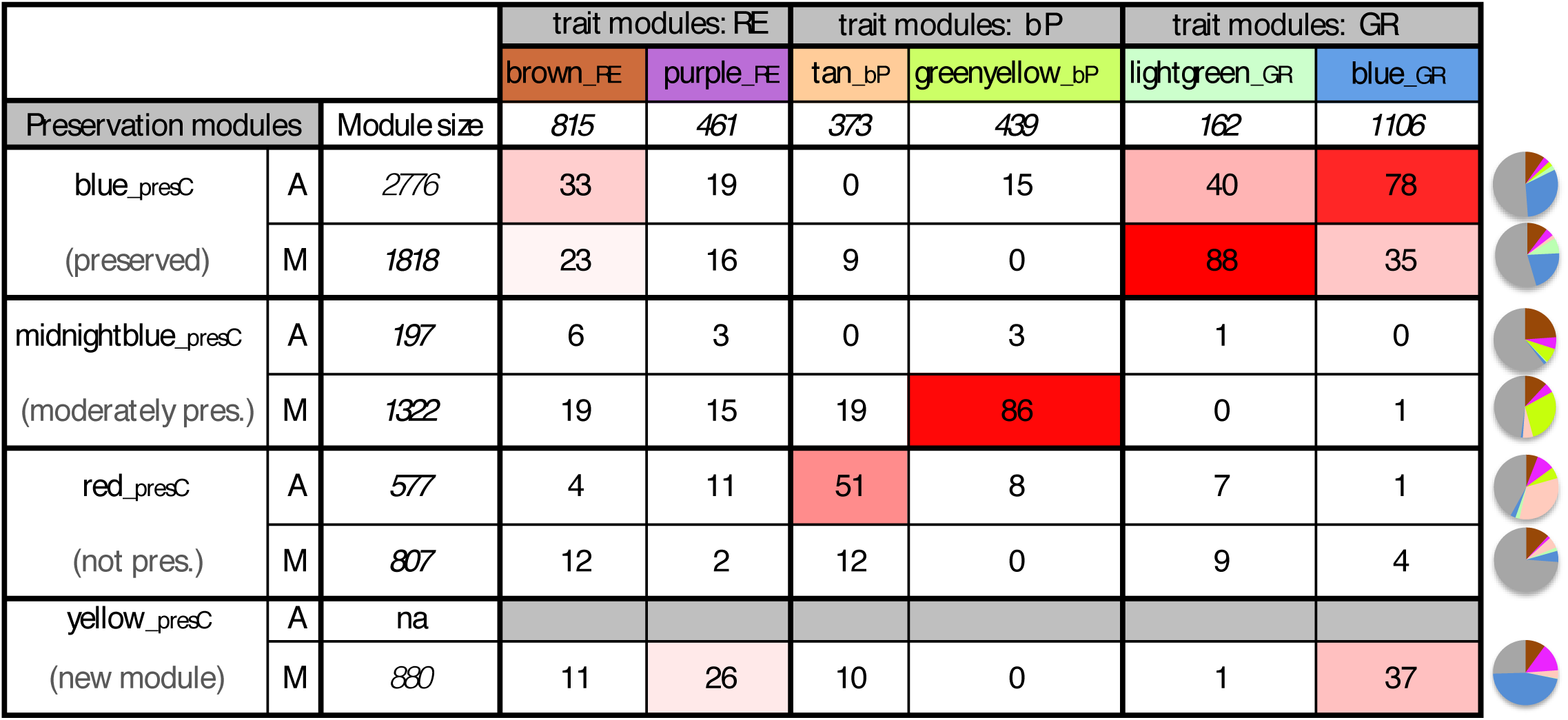
Association between trait-associated network and preservation network modules. The heatmap lists percentages of genes shared between trait-associated modules (columns) and ancient and modern preservation network modules (rows), i.e. percentages of genes both present in a given trait-module and a given preservation module. Numbers in italics represent module size (gene number) of trait-associated and preservation modules. Pie charts show the fractions of each of the six trait modules overlapping with a given preservation module. The grey fraction represents preservation module genes not overlapping with any of the trait modules. *Note: due to separate construction of the trait-associated network and the preserved networks, shared colours of modules in these networks (trait-associated and preservation networks) do not imply that modules are identical.*

**Fig. 5.**
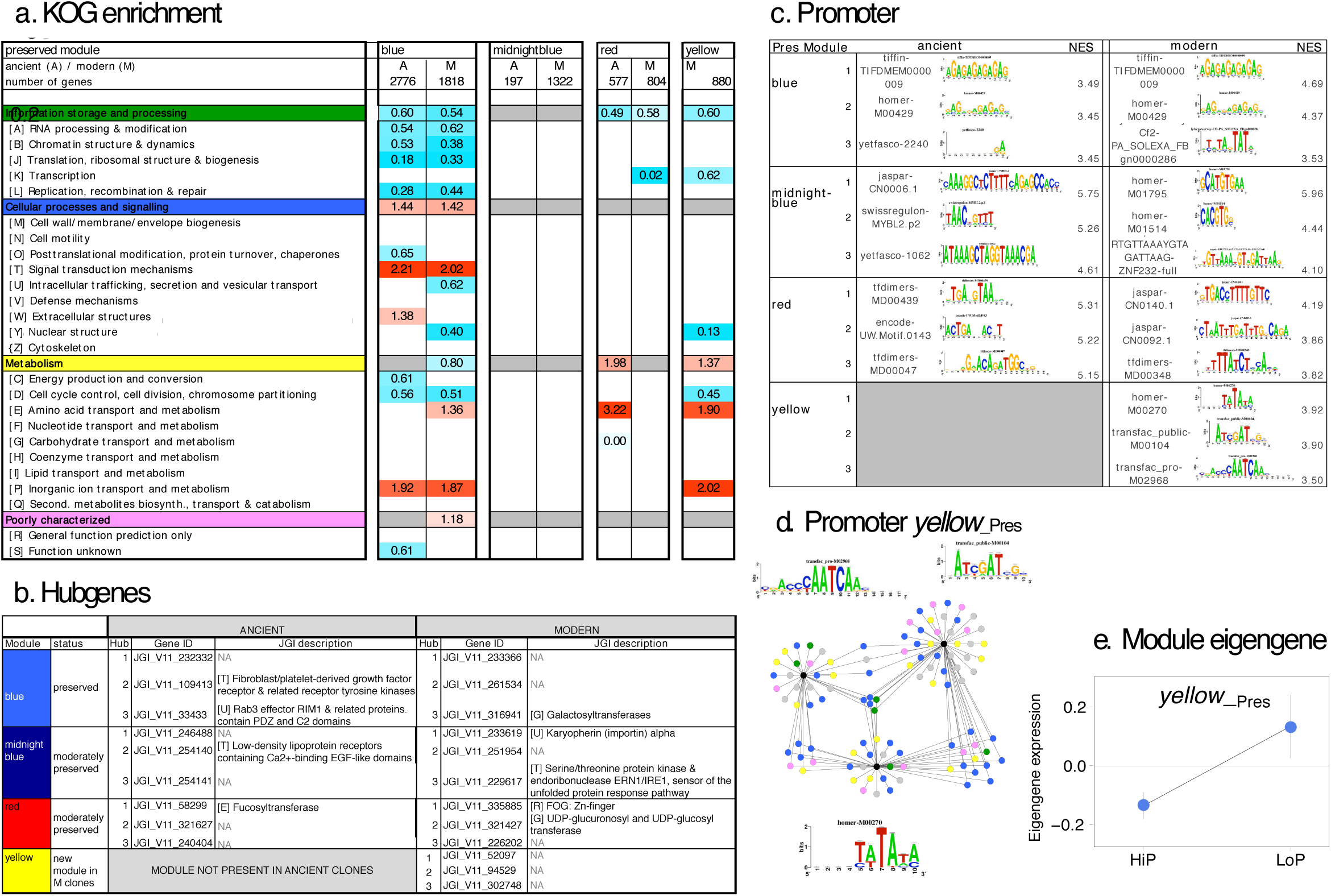
Functional analysis of the preservation modules represented in trait-associated modules as shown in Fig. 4. a. KOG enrichment of the top preserved or divergent modules identified in the preservation network analysis. Enrichment scores (odds-ratios, FDR <0.05) indicating over- and under-representation of genes in defined functional groups (colour-coded as red and blue, respectively) are calculated separately for each module and can therefore only be directly compared within columns. b. Top three hubgenes of the preserved/divergent modules listed in A. c. Promoter motif analysis of the preserved/divergent modules listed in A. NES= normalised enrichment score. d. Cytoscape network of potential promoter target genes. Colour coding (i.e., green, blue, yellow, pink and grey) indicates functional association of nodes to distinct KOG categories (for details see insert in Fig. 3). Black nodes represent promoter motif binding sites (also shown as sequence logo). e. Eigengene expression of the *yellow*__Pres_ module in high P (HiP) and low P (LoP) conditions.

Strikingly, our preservation analysis identified a new module (*yellow*_Pres,_ Fig. S6c, S9) in the modern *Daphnia* clones that contained a set of 880 co-expressed genes and lacked a comparable modular structure in the ancient gene network. Such new modules provide evidence for evolutionary novelty on the level of transcription^33^. All genes of this module were more highly expressed in P-limited conditions, thus revealing a high degree of gene expression plasticity (Fig. 5e, S9).

#### Association between preservation network and trait-associated network modules

In order to evaluate links between phenotypic divergence, gene co-expression and network preservation, we focussed further analysis on a subset of preservation modules that each comprised a large fraction of the trait-associated module genes (Figs. 4 and S10). These represent four preservation levels: preserved (*blue*_Pres_), moderately preserved (*midnightblue*_Pres_), non-preserved (*red*_Pres_) and a newly formed, modern module (*yellow*_Pres_). Together, the sum of genes of the six trait-modules constituted ∼50% or more of either an ancient or a modern counterpart of these preservation module pairs, or the new *yellow*_Pres_ module (pie charts in Fig. 4). The presence of trait-module genes was particularly pronounced in *yellow*_Pres_ which contained ∼75% of trait-associated genes.

Functionally, the *blue*_Pres_ module was similar in ancient and modern clones (Fig. 5a), with enrichments in signal transduction and metabolic processes involved in inorganic ion transport. As expected, the moderately preserved (*midnightblue*_Pres_) and non-preserved (*red*_Pres_) modules did not show such functional similarities. The analysis of promoter motif enrichment supported these findings for preservation and divergence with largely identical enrichment of promoter motifs in the preserved *blue*_Pres_, but distinct sets of promoter binding sites in the moderately and non-preserved modules (Fig. 5c). In contrast, hubgenes differed between ancient and modern modules regardless of their preservation status (Fig. 5b). These hubs covered a range of cellular processes, including signal transduction mechanisms and amino acid and carbohydrate transport. Over 50% of the top three hubgenes had no functional annotation.

The largest fraction of overlap between trait-modules and *blue*_Pres_ was observed with the GR-modules *lightgreen*__GR_ and *blue*__GR_ (Fig. 4), suggesting that these modules are largely unaffected by evolutionary change. This relationship indicates shared molecular processes in ancient and modern clones that drive this phenotypic trait, which is supported by an almost identical functional enrichment (Fig. 3a, 5a). To a smaller degree, *brown*__RE_ genes were also shared with the *blue*_Pres_ modules (Fig. 4), indicating that the molecular drivers of RE are at least partly preserved between ancient and modern *Daphnia.*

The moderately preserved *midnightblue*_Pres_ and the non-preserved *red*_Pres_ had the largest overlap with the bP-associated modules *greenyellow*__bP_ and *tan*__bP._ Interestingly, this overlap was observed only with one of the module counterparts of *midnightblue*_Pres_ and *red*_Pres_: the trait module *greenyellow*__bP_ shared 86% of its genes, and one promoter motif with the modern *midnightblue*_Pres_ (Fig. 3c, 4 and 5c). *Tan*__bP_ shared about 50% of its genes, their functional enrichment, and its gene regulatory network (i.e., shared promoter binding sites and top hub genes, Fig. 3-5) with ancient *red*_Pres_. These pattern suggest that bP is driven by divergent molecular processes in ancient and modern clones.

*Yellow*_Pres_, the new modern module, contained about 75% of the six trait-associated modules (Fig. 4, pie charts). A large portion of these genes were found in *blue*__GR_ which shared 37% of its genes with *yellow*_Pres_, and *purple*__RE_ which shared 26% of its genes with *yellow*_Pres_. This overlap with the novel *yellow*_Pres_ module suggests that both of these trait-modules include a large fraction of genes with an evolutionarily altered expression pattern. *Yellow*_Pres_ combined functionally enriched groups of *purple*__RE_ (amino acid transport and metabolism) and *blue*__GR_ (inorganic ion transport and metabolisms) (Fig. 3a and 5a). The *yellow*_Pres_ gene regulatory network (Fig. 5d) resembled that of the *blue*__GR_ module (Fig. 3c) with many genes potentially co-regulated by at least two transcription factors.

## Discussion

Eutrophication, with P as a major pollutant, is a pervasive and fundamental threat to aquatic ecosystems of our planet. In order to preserve the integrity of these ecosystems, we need a detailed understanding of the molecular mechanisms that allow keystone species to adapt to and persist in altered environments, facilitating the maintenance of their vital ecological services.

To advance this understanding, we linked gene co-expression networks with changes in phenotypic traits using resurrected *Daphnia* isolates separated by centuries of evolution and anthropogenic change. Network analyses allowed us to identify gene clusters and their networks that underlie organismal responses to environmental variation. To provide a direct phenotype-genotype link, we applied such a network approach and combined it with quantitative trait data observed in members of a single *Daphnia* population before and after a historic shift in nutrient supply associated with modern agricultural activities^14,17^. Specifically, we explored the transcriptional regulation of two physiological traits related to P acquisition (retention efficiency (RE), somatic P content (bP)), and a higher-order phenotypic trait dependent on RE and bP, i.e. somatic growth rate (GR) using a trait-associated gene co-expression network. The resulting network suggests a strong relationship of transcriptional responses with P-supply, with over 50% of the 17 observed modules significantly associated with the P-related phenotypic traits.

Complementing the trait-associated network, the use of network preservation statistics can identify the ‘wiring’ of molecular mechanisms that are shared or divergent between ancestral isolates and their modern counterparts^33^. Our results highlight a highly preserved network structure with >70% of the ancient *Daphnia* modules preserved in modern descendants that was also reflected by shared gene regulatory mechanisms in ancient and modern modules (i.e. *blue*_Pres_). Such a pattern is not unexpected in members of the same population, considering a similar level of preservation in closely related taxa that diverged from a common ancestor several million years ago (such as humans and chimpanzees^33^). However, the analysis of network preservation also revealed patterns of evolutionary divergence of ancient *Daphnia* and their modern descendants in individual modules.

Adaptation to environmental change is typically associated with divergent gene expression patterns^20,22,29^. The trait-associated, transcriptional responses of ancient and modern *Daphnia* observed here support such findings (Fig. 2b), providing evidence of distinct evolved patterns of gene expression: conserved gene expression plasticity (*brown*__RE_, *lightgreen*__GR,_ *blue*__GR_), genetic assimilation (tan_*_bP*_) and evolved plasticity (*purple*__RE,_ *greenyellow*__bP_), *sensu* Renn and Schumer^21^. Given that the interaction of genes within modules is stronger than between modules, and that modules are regarded as ‘semi-independent’ units that evolve independently due to reduced pleiotropic constraints^53-55^, the coexistence of these observed gene expression patterns within a single network strongly supports the idea of individual evolutionary trajectories of these trait-associated modules.

Plastic gene expression in response to P-supply was common to all focal modules, but was not limited to ancient or modern clones (Fig. 2, Table 1). For example, RE correlated strongly with modules that showed signs of newly evolved plasticity (i.e., *purple*__RE_). In contrast, genes in both GR-associated modules (i.e., *blue*__GR_, *lightgreen*__GR_) maintained similar gene expression plasticity in ancient and modern clones. While our data suggest that gene expression is often plastic, such plasticity did not always translate into similar plasticity in the tested phenotypic traits (e.g., GR): a plastic gene expression in ancient clones was less obvious in their phenotypic response. A potential explanation for this may be the complexity of this trait that depends on many other factors, including nutrient availability and assimilation, and other factors involved in cellular and developmental processes.

Phenotypic plasticity can have many forms, including active plasticity (actively regulated by an organism, e.g. by regulatory genes, often thought to be adaptive) and passive plasticity (often considered non-adaptive) which Whitman and Agrawal^56^ refer to as “simple susceptibilities to chemical and physical stresses“. However, it is likely that these forms of plasticity are not mutually exclusive. Indeed, our data underline that the distinction between these two types of plasticity is not straightforward. Although the network preservation analysis alone could not differentiate between these types of plasticity, its results indicate evolutionary novel patterns of gene expression in the form of a new module (*yellow*_Pres_) that was unique to modern *Daphnia* clones. By integrating preservation and trait-associated networks, we were able to establish a link between the evolution of gene expression and phenotypic plasticity. The presence of a high percentage of the *yellow*_Pres_ genes in two trait-associated modules, *purple_*_RE_ (‘adaptive plasticity’) and *blue*_*_*GR_ (‘non-adaptive plasticity’), indicate the coexistence of different types of plasticity in a single module (here: *yellow*_Pres_), suggesting that different forms of plasticity may coexist within single traits.

### Concluding remarks

To genuinely advance the understanding of phenotypic evolution, comprehensive methods are required that consider entire organisms instead of single traits^57^. Such a holistic understanding is vital in order to predict evolutionary trajectories that result from major geochemical shifts that currently affect our planet, and are essential for the development of conservation strategies. The results presented here are a contribution towards such an understanding, and emphasise the need for an integrative approach that combines physiological and ‘omics data in keystone species.

Our study highlights the power of the applied strategy and motivates future research in several areas:

(1) Genomic manifestation: theory predicts relaxed selection on loci underlying genetically controlled phenotypic plasticity and thus higher genomic variation in associated genes^53^. This raises the question whether the observed differences in gene expression between ancient and modern *Daphnia* clones are manifested in the genomic sequence, for instance as increased genetic divergence in modules with evolved plasticity.

(2) Molecular regulators of phenotypic plasticity: while transcriptional regulation provides a critical mechanism for organisms to respond rapidly and efficiently to environmental change^58^, the contribution of different molecular regulators of plasticity (*e.g.,* epigenetic modifications, transcription factors) remain largely unknown. Our work specifically points to the importance of histone modifications in this context.

(3) Role of hubgenes: on a functional level, recent advantages in molecular techniques now allow for a detailed testing of network structures to confirm if molecular cascades collapse when predicted hub-genes are modified via gene editing approaches (e.g., RNAi or CRISPr and Talen techniques^59-61^).

Contrasting resurrected members of populations that lived hundreds of years ago with their modern descendants, as done here, is a rare opportunity to track evolutionary trajectories in natural environments. Our study highlights the prospects of resurrection ecology when integrated with modern biology. It further emphasises the applicability of this approach to numerous other organisms that produce dormant stages with long-term viability, and its significance for an in-depth understanding of evolutionary adaptation to global environmental change.

## Author contribution

All authors contributed equally to this manuscript.

## Supplementary Figure and Tables

Fig. S1. Hierarchical clustering tree of the trait-associated modules using module eigengenes.

Fig. S2. Heatmap showing correlation coefficients and p-values of the relationship between module eigengenes of the 17 trait-associated modules and phenotypic traits: P retention efficiency (RE), body P concentration (bP), and somatic growth rate (GR).

Fig. S3. Gene significance (for details see Horvath^23^) across modules and phenotypic traits. Module colour coding as in Fig. S2.

Fig. S4. Functional enrichment (KOG) of trait-associated modules that were significantly positively (yellow highlight) or negatively (green highlight) correlated with one of the traits (R^2^> 0.4 and p ≤ 0.005). Enrichment scores (odds-ratios, FDR <0.05) indicating over- and under-representation of genes in defined functional groups (colour-coded as red and blue, respectively) are calculated separately for each module and can therefore only be directly compared within columns.

Fig. S5. a-f: Circular plots of gene expression patterns in six trait-associated modules. g-m (inserts of a-f): Circular plots of expression patterns of genes shared between trait modules and preservation modules of modern clones (see also Fig. 4).

Tracks of the graphs show (from outside to inside): *Track 1*- KOG annotation; letters correspond to those of KOG enrichment tables (i.e., Figs. 3, 5, S4, and S8). *Track 2* - Normalised log_2_-transformed intensities. *Track 3 -* Total connectivity (orange bars) and within connectivity (yellow bars) of module members (genes). *Track 4 -* Gene expression patterns (log_2_ ratios of normalised gene intensities) of ‘within-treatment’ ratios (dark green bars - A:M ratio at high P (HiP) food; light green bars - A:M at low P (LoP) food). Positive lightgreen or darkgreen bars of ‘within-treatment’ ratios indicate higher expression in ancient clones, negative lightgreen or darkgreen bars indicate higher expression in modern clones. *Track 5 -* Gene expression pattern (log_2_ ratios of normalised gene intensities) of ‘within-era’ (ancient and modern) ratios (pink bars - LoP:HiP in ancient clones; blue bars - LoP:HiP in modern clones). Positive blue or pink bars of ‘within-era’ ratios indicate higher expression under low P food, while negative blue or pink bars indicate higher expression under high P food. Innermost track indicates the name and colour of modules shown.

Fig. S6. Preservation networks.

a. Gene dendrogram of ancient network resulting from average linkage hierarchical clustering with module annotation (colour bar). Red lines indicate cut-off used for module construction.

b. Gene dendrogram of modern network resulting from average linkage hierarchical clustering with module annotation (colour bar).

c. Cross-tabulation of genes associated with ancient modules (rows) and modern modules (columns). Numbers next to the module colours indicate module size (total number of genes in each module), while numbers within the table represent number of genes overlapping (see colour-coded scale on the right) between respective ancient and modern modules with calculated p-statistics.

Fig. S7. Median Rank and Z-summary statistics indicating preservation status of modules in ancient and modern *Daphnia* clones. Lower Median Rank values and higher Z-summary values indicate higher preservation, respectively (for details see Langfelder and Horvath^32^).

Fig. S8. Functional enrichment (KOG) for the most highly preserved and the non-preserved modules of the preservation networks (detailed in Fig. S7). Enrichment scores (odds-ratios, FDR <0.05) indicating over- and under-representation of genes in defined functional groups (colour-coded as red and blue, respectively) are calculated separately for each module and can therefore only be directly compared within columns.

Fig. S9. Circular plot of gene expression patterns in the *yellow*__Pres_ module that was newly formed in the modern preservation network. Colour coding in all tracks as in Fig. S5. Tracks of the graphs show (from outside to inside): *Track 1*- KOG annotation. *Track 2* – blue and purple dots indicate genes shared between *yellow*__Pres_ and *blue*__GR_ or *purple*__RE_, respectively. *Track 3* - Normalised log_2_-transformed intensities. *Track 4 -* Total connectivity (orange bars) and within connectivity (yellow bars) of module members (genes). *Track 5 -* Gene expression pattern (log_2_ ratios of normalised gene intensities, LoP:HiP) of modern clones. Positive blue bars indicate higher expression under low P food, while negative blue bars indicate higher expression under high P food.

Fig. S10. Gene overlap between the six trait modules most highly associated with one of the phenotypic traits (RE, bP and GR). Bars represent the fraction of the respective trait-module that was shared with one of the preservation modules listed along the x-axis.

Table S1. Network statistics of the trait-associated network.

Table S2. Network statistics of the preservation networks. Highlighted modules of the ancient network denote modules that were absent in the modern network and vice versa.

## Methods

### Phosphorus-related phenotypic traits

Our study capitalised on previously collected data that provide phosphorus-related phenotypic data (phosphorus retention efficiency (RE), somatic growth rate (GR)^14^; somatic P concentration (body P, or bP)^30^), and gene expression data of ancient and modern *Daphnia pulicaria* genotypes in response to distinctive phosphorus regimes^17^. Briefly, while RE measures the amount of P retained by neonate *Daphnia* from dietary sources over a short period of time (12 h), bP represents the amount of P retained over five days)^30^.

In order to obtain this data, resurrected genotypes of the cyclical parthenogen *D. pulicaria* were used. Specifically, *Daphnia* genotypes included two ancient genotypes with an estimated age of **∼**600**–**700 years (A1, A2) and their modern descendants, two genotypes with an estimated age of 5**–**10 years (M1, M2). The ancient and modern genotypes evolved in contrasting P regimes (ancient = low P environment, modern = high P environment (for details see^14^). Significant phenotypic shifts were observed in P-kinetics between these ancient and modern genotypes, with the former retaining P more efficiently and showing higher growth rates at low P conditions than the latter. Therefore, these four genotypes were ideal to explore transcriptional patterns underlying diverging phenotypic responses to cultural eutrophication.

To test both physiological and transcriptional responses, animals were exposed to high and low P food conditions (*Scenedesmus acutus*; high P = C:P ratio of ∼150, low P = C:P ratio of ∼750) at a C concentration of 3 mg C L^-1^ day^-1^. Gene expression data were obtained using a NimbleGen 12-plex long oligonucleotide microarray (GEO: GPL11278)^62^. For details of RNA hybridisation and normalisation of raw intensities see Row Chowdhury et al. 2015^17^.

### Weighted gene co-expression network analysis (WGCNA)

We used the R package WGCNA for weighted correlation network analysis^32^ to construct gene co-expression networks. The general procedure of WGCNA has been described in detail elsewhere^63^. In brief, we constructed a signed overall co-expression network using gene expression data^17^ that assessed transcriptional responses of ancient and modern daphnids exposed to contrasting P regimes. Specifically, we used normalized intensities as input, and limited our analyses to genes that were identified as differentially expressed in one of the tested conditions (FDR < 0.05), using *limma*^64^ analysis (for details see^17^). First, we calculated a similarity co-expression matrix (Pearson’s correlation) for all genes, and transformed these values to an adjacency matrix by using the soft thresholding power (beta = 10), to which co-expression similarity was raised. This step aims to emphasize and reduce the contribution of strong and weak correlations, respectively, on an exponential scale, resulting in a network exhibiting approximate scale-free topology. Next, all genes were hierarchically clustered based on a dissimilarity measure of topological overlap which measures interconnectedness for a pair of genes^63^. The resulting gene dendrogram was used for module detection with the Dynamic Tree Cut method (minimum module size = 100, cutting height = 0.2). Gene modules corresponding to the branches cut-off of the gene tree were colour-coded. WGCNA provides a feature to control for genes that cannot be associated with one of the identified expression patterns (the “grey” module), and therefore cannot be placed into any of the existing modules. However, the absence of a grey module in our analysis indicated clear expression patterns throughout the entire dataset. WGCNA further allows for summarizing obtained modules by using eigengenes, which are defined as the first principal component of the expression matrix for each module. Network metrics for individual genes were calculated, including gene significance (i.e., the absolute value of the trait-gene correlation) and module membership (i.e., the module eigengene-gene correlation). Module membership was used to define the top central genes for modules of interest (i.e., hub genes). To assess the contribution of sets of co-expressed genes (i.e., modules), gene expression data was correlated with phenotypic trait data (see above) using the plotDendroAndColors function^32^. Here, we focus on the two most highly correlated modules per phenotypic trait with a module-trait correlation R^2^ ≥ |0.4| and a statistical significance of p ≤ 0.005 (for a complete list of trait-module correlations see Fig. S2).

### Preservation network analysis

Module preservation statistics were computed using the modulePreservation function (5000 permutations) implemented in WGCNA^52^. Network module preservation statistics quantify how density and connectivity patterns of modules defined in a reference data set (here: ancient genotypes) are preserved in a test data set (here: modern genotypes). Network adjacency comparisons are superior estimates of module preservation to standard cross-tabulation techniques. We therefore used network adjacency comparison to assess preservation of gene co-expression patterns in ancient and modern modules. For this, we constructed two networks separately for ancient and modern *Daphnia* clones. The overall significance of the preservation was assessed using Zsummary and median rank statistics^52^. Based on the thresholds proposed by Langfelder et al.^52^, Zsummary < 2 indicates no preservation, 2 < x < 10 weak to moderate evidence of preservation, and >10 strong evidence of module preservation across networks. Median rank statistics further consider module size for preservation assessment^52^.

### Connection between trait-associated and preservation networks/ modules

In order to assess links between phenotypic divergence, gene-coexpresssion and network preservation, we traced the gene sets of the six focal phenotype modules in modules of the ancient and modern preservation networks. For this, we identified the percentage of genes of a given phenotype module that was present in the modules of the preservation networks (Fig. S6). For closer examination, we chose preservation modules (either the ancient or the modern module of a given colour, or the new yellow module) that contained at least 1/5th of the genes of a phenotype module. Statistical analyses were performed with the R Statistical Software, version 3.4.1 (R Core Team 2017)^65^.

### Functional enrichment analyses

All predicted protein-coding gene models were functionally assigned to orthologous groups (euKaryotic Orthologous Groups, KOG)^66^ as defined by the Joint Genome Institute ^67^. Fisher’s exact tests (p ≤ 0.05) were applied to assess enrichment of up- or down-regulated genes in the different KOG categories. The reference set for statistical testing comprised all genes that entered the network analysis.

### Promoter motif analysis

To describe gene regulatory patterns we identified the three most enriched promoter motifs for each phenotype module. We ran a promoter motif analysis with the web-based *Daphnia* cis-target (http://daphniacistarget.aertslab.org) following Spanier et al^16^. For the trait-associated network, we used as input the complete gene sets of the six modules most highly correlated with the phenotypic traits RE (*purple*__RE_, *brown*__RE_), bP (*tan*_*_*bP_, *greenyellow*__bP_) and GR (*blue*__GR_, *lightgreen*__GR_). For the preservation networks, we used the gene sets of the three module pairs discussed in this work (ancient and modern *blue*_Pres_, *midnightblue*_Pres,_ *red*_Pres)_, and the modern *yellow*_Pres_ module. The promoter analysis was run with default settings and a normalised enrichment score (NES) of 2.5, using the database containing motifs 300bp upstream of transcription start sites.

### Multivariate analysis of gene expression profiles

Multidimensional scaling (MDS) was used to plot the gene expression profiles of ancient and modern clones in response to high and low P food using the isoMDS() function in the MASS package version 7.3-51.1^68^ for the R Statistical Software, version 3.4.1^65^. Normalised intensities were used to calculate Euclidian distances between all samples.

### Linear mixed effects models

To test the linear effects of evolutionary history and phosphorus supply on module eigengene expression, we fitted nested linear mixed effects models with the ‘lme4’ package^69^ for the R Statistical Software, version 3.4.1^65^. The dependent variable Module Eigengene (ME) expression was transformed prior to the analysis in order to met the requirements of the linear model. For this, a Box-Cox transformation was performed using the boxcox() function from the MASS package, version 7.3-51.1^68^. Because the transformation requires positive numbers, the value 1 was added to all ME expression values. Models were fitted for the fixed effects of ‘age’ (ancient or modern) and treatment (high or low P food) and their interaction by maximum likelihood (with the ‘REML = FALSE’ argument in the model specification), and the random factor ‘clone’ (4 clones, two modern and two ancient), nested within ‘age’, and compared against a null model that excluded all fixed effects using a likelihood ratio test.

### Visualizations

Cytoscape v3.6.1^70^, a software environment for integrated models of bimolecular interaction network was employed to visualise network structure of the six most highly correlated trait-associated modules found in our analysis. The input for cytoscape networks were exported from WGCNA, using the function exportNetworkToCytoscape () of the WGCNA package. Specifically, the cytoscape network was created including only nodes that are connected by edges (intramodular connectivity k) with a weight >0.4, thus reducing complexity of the graph.

Cytoscape v3.6.1 was also used to visualise networks of promoter motif target genes.

Circular plots were created using the *circlize* package version 0.4.5^71^ for the R Statistical Software, version 3.4.1^65^.

## Acknowledgements

We are grateful to Priyanka Roy Chowdhury, Katja Pulkkinen and Andrew Beckerman for valuable discussions of the manuscript. Funding for generation of transcriptome and phenotypic data was provided by the US National Science Foundation collaborative grants #0924289, #1256881 and #0924401. DB was funded by independent research fellowships from the German Research Foundation (BE5288/2-1, BE5288/3-1). DF received funding from the European Union’s Horizon 2020 research and innovation programme under the Marie Skłodowska-Curie grant agreement No. 658714. MW received funding from the European Union’s Intra-European Fellowships for Career Development (FP7-MC-IEF), Marie Sklodowska-Curie grant agreement No. 629892 and from the Norwegian Research Council, IS-TOPP No. 238820.

