## Supplementary Material for "Dissecting the transcriptomic basis of phenotypic evolution in an aquatic keystone grazer"

Figure S1

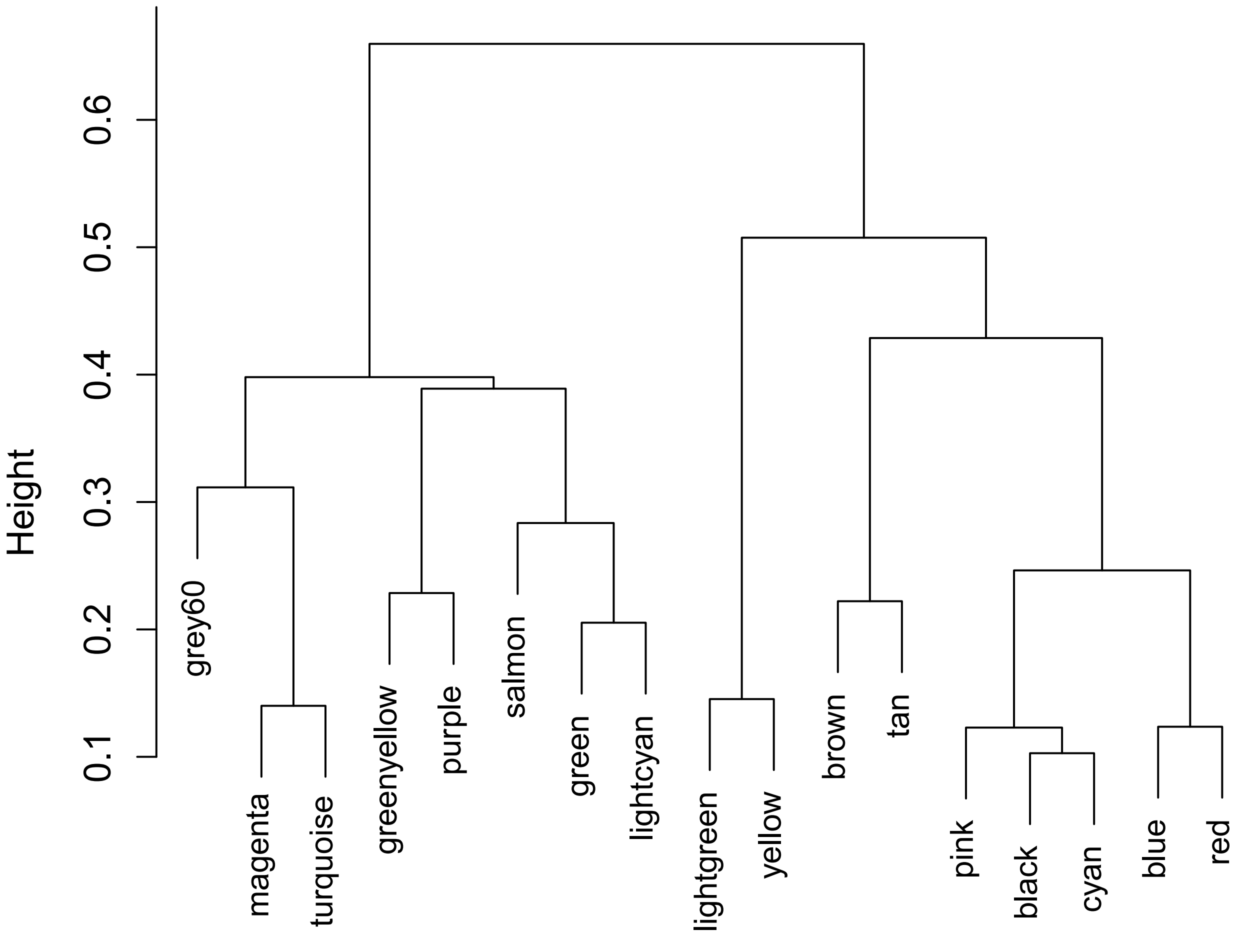
Height
0.1
0.2
0.3
0.4
0.5
0.6
grey60
magenta
turquoise
greenyellow
purple
salmon
green
lightcyan
lightgreen
yellow
brown
tan
pink
black
cyan
blue
red

Figure S2

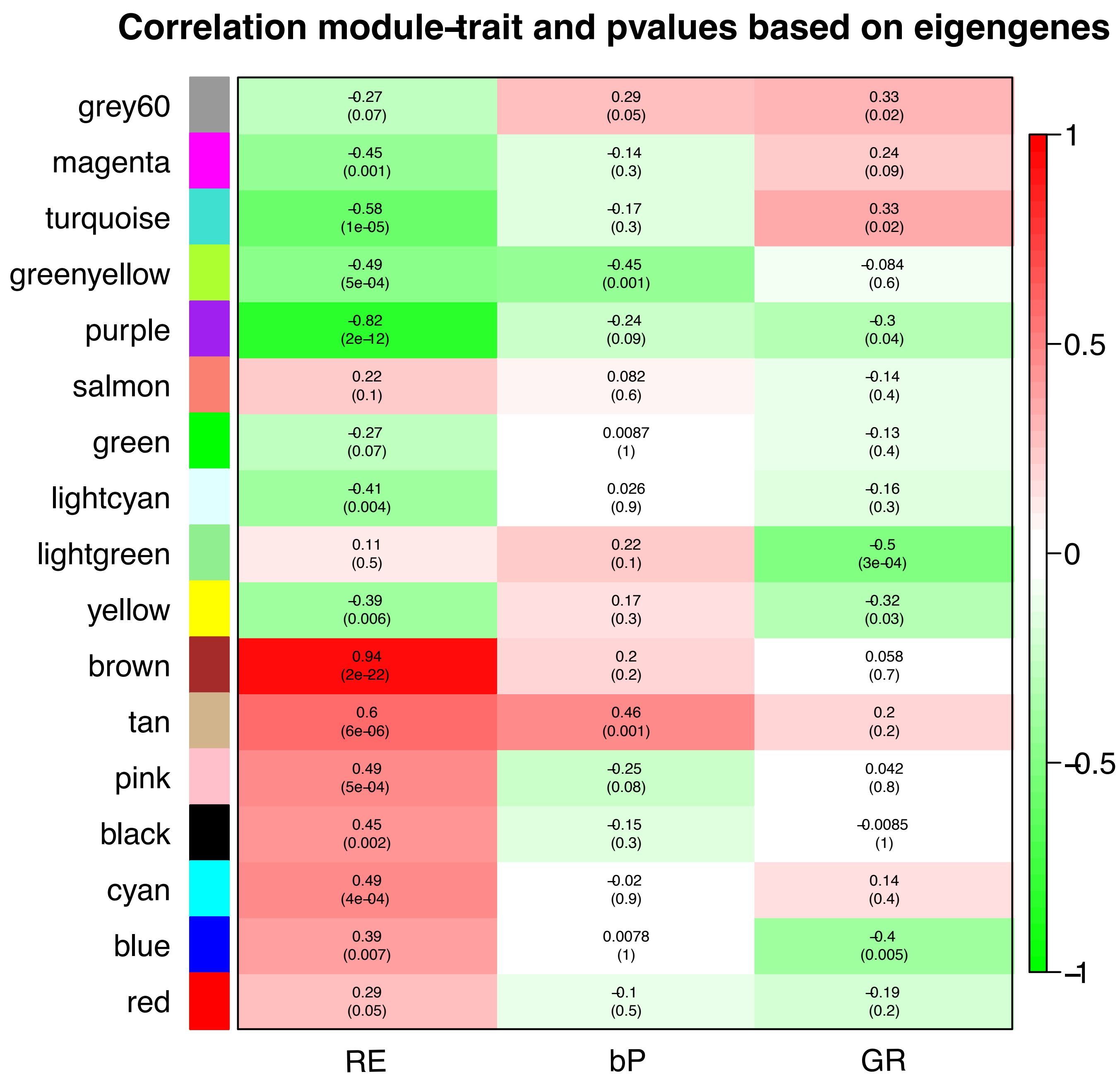

Correlation module-trait and pvalues based on eigengenes
grey60
magenta
turquoise
greenyellow
purple
salmon
green
lightcyan
lightgreen
yellow
brown
tan
pink
black
cyan
blue
red
-0.27 (0.07)
0.29 (0.05)
0.33 (0.02)
-0.45 (0.001)
-0.14 (0.3)
0.24 (0.09)
-0.58 (1e-05)
-0.17 (0.3)
0.33 (0.02)
-0.49 (5e-04)
-0.45 (0.001)
-0.084 (0.6)
-0.82 (2e-12)
-0.24 (0.09)
-0.3 (0.04)
0.22 (0.1)
0.082 (0.6)
-0.14 (0.4)
-0.27 (0.07)
0.0087 (1)
-0.13 (0.4)
-0.41 (0.004)
0.026 (0.9)
-0.16 (0.3)
0.11 (0.5)
0.22 (0.1)
-0.5 (3e-04)
-0.39 (0.006)
0.17 (0.3)
-0.32 (0.03)
0.94 (2e-22)
0.2 (0.2)
0.058 (0.7)
0.6 (6e-06)
0.46 (0.001)
0.2 (0.2)
0.49 (5e-04)
-0.25 (0.08)
0.042 (0.8)
0.45 (0.002)
-0.15 (0.3)
-0.0085 (1)
0.49 (4e-04)
-0.02 (0.9)
0.14 (0.4)
0.39 (0.007)
0.0078 (1)
-0.4 (0.005)
0.29 (0.05)
-0.1 (0.5)
-0.19 (0.2)
RE
bP
GR
1
0.5
0
-0.5
-1

### Figure S3

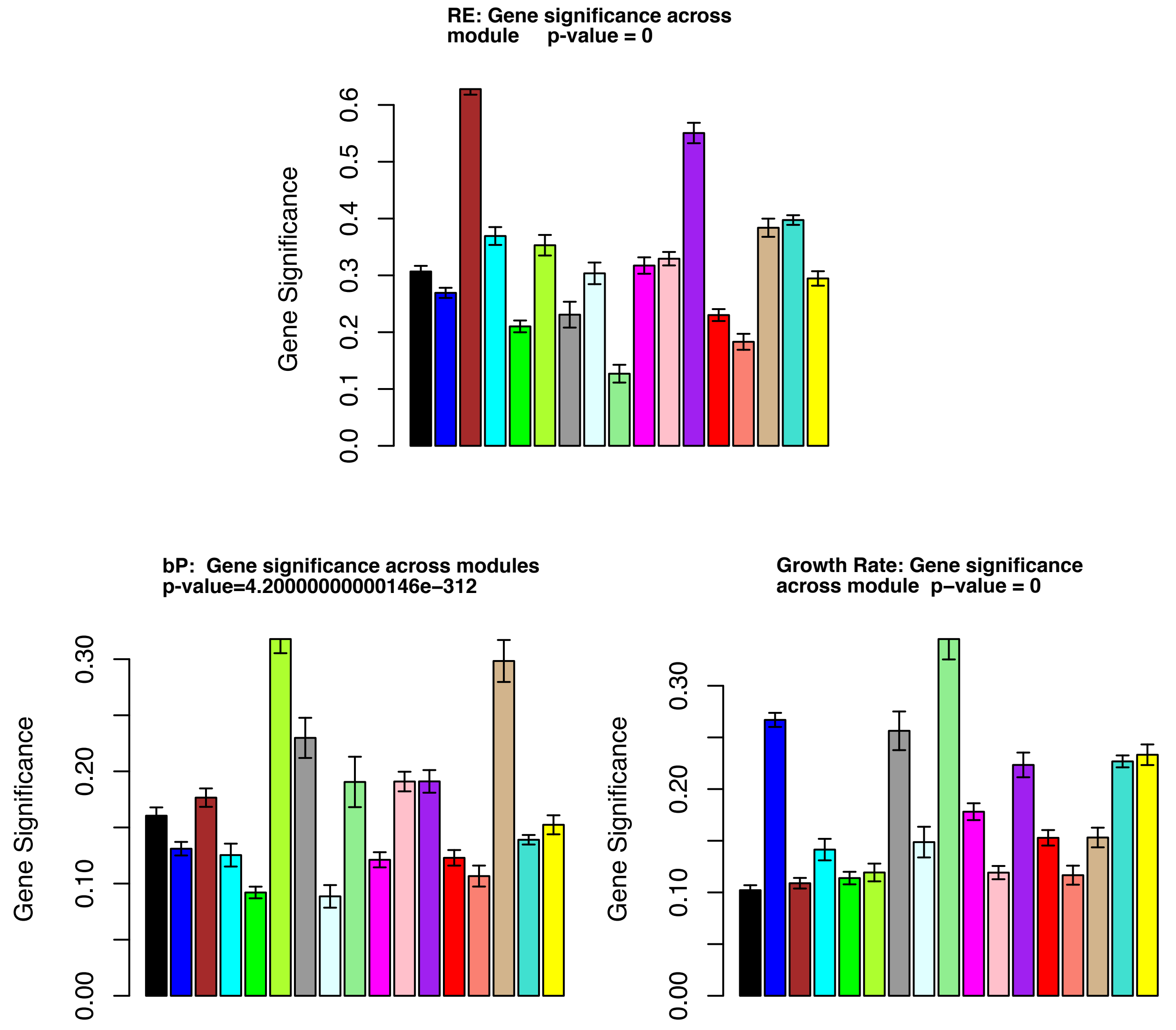
RE: Gene significance across
module p-value = 0
Gene Significance
0.0
0.1
0.2
0.3
0.4
0.5
0.6
bP: Gene significance across modules
p-value=4.20000000000146e-312
Gene Significance
0.00
0.10
0.20
0.30
Growth Rate: Gene significance
across module p-value = 0
Gene Significance
0.00
0.10
0.20
0.30

### Figure S4

| Phenotype | RE | | | | | | | | | | GR | | bP | |
|---|---|---|---|---|---|---|---|---|---|---|---|---|---|---|
| Module colour | brown | tan | pink | cyan | black | purple | green-yellow | magenta | tuquoise | lightcyan | light-green | blue | tan | green-yellow |
| module-trait (RE) correlation (p-value) | 0.94 (2e-22) | 0.60 (6e-06) | 0.49 (5e-04) | 0.49 (0.0004) | 0.45 (0.002) | -0.82 (2e-12) | -0.49 (0.0005) | -0.45 (0.001) | -0.58 (0.00001) | -0.41 (0.004) | -0.5 (0.0003) | -0.4 (0.005) | 0.46 (0.001) | -0.45 (0.001) |
| # genes | | | | | | | | | | | | | | |
| Information storage and processing | 0.45 | 0.58 | 0.56 | 0.44 | 0.47 | | | 1.84 | 1.38 | 0.45 | | 0.50 | 0.58 | |
| [A] RNA processing & modification | | | | | | | | | | | | 0.33 | | |
| [B] Chromatin structure & dynamics | | | | | | | | | 1.82 | | | | | |
| [J] Translation, ribosomal structure & biogenesis | | | | | 0.28 | | | 2.31 | 0.45 | | | 0.23 | | |
| [K] Transcription | | | | | | | | 1.81 | 1.48 | | | | | |
| [L] Replication, recombination & repair | | | | 0.00 | 0.19 | | | | 1.86 | | | 0.28 | | |
| Cellular processes and signalling | 1.41 | | | | 1.27 | 0.75 | | | 0.69 | | | 1.43 | | |
| [M] Cell wall/membrane/envelope biogenesis | | | | | | | | | 0.21 | | | | | |
| [N] Cell motility | | | | | 6.10 | | | | | | | | | |
| [T] Signal transduction mechanisms | | | | | 1.33 | | | | 0.69 | | | 1.99 | | |
| [U] Intracellular trafficking secretion and vesicular transport | | | | | | | | | | | | | | |
| [V] Defense mechanisms | | | | 2.42 | | | | | 0.53 | | | | | |
| [W] Extracellular structures | | | | 2.49 | 2.86 | | | | 0.31 | | | | | |
| [Y] Nuclear structure | | | | | | | | | | | | | | |
| Metabolism | | 1.60 | 1.50 | 2.05 | 1.39 | 1.70 | 1.58 | | | 3.16 | 1.85 | | 1.60 | 1.58 |
| [C] Energy production and conversion | | | | | | | | | 0.41 | | | | | |
| [D] Cell cycle control, cell division, chromosome partitioning | | | 0.27 | 0.12 | 0.32 | | | 2.13 | 3.29 | | | 0.55 | | |
| [E] Amino acid transport and metabolism | | 2.18 | | | 0.60 | 2.55 | 2.29 | | 0.69 | 3.96 | 3.89 | | 2.18 | 2.29 |
| [F] Nucleotide transport and metabolism | | | | | | | | | | | | | | |
| [G] Carbohydrate transport and metabolism | | | | | 1.51 | | | | | 4.79 | | 0.46 | | |
| [H] Coenzyme transport and metabolism | | | | | | | | | | | | | | |
| [I] Lipid transport and metabolism | | | 2.07 | 4.78 | 3.32 | | | | | | | | | |
| [P] Inorganic ion transport and metabolism | | | | | | | | | | | | 2.08 | | |
| [Q] Second. metabolites biosynth., transport & catabolism | | | | | 2.78 | | | | | | | | | |
| Poorly characterized | | | | | | | | | | 0.48 | | | | |
| [R] General function prediction only | | | | | | | | | | 0.44 | | | | |
| [S] Function unknown | | | | 0.26 | | | | | | | | | | |

### Figure S5

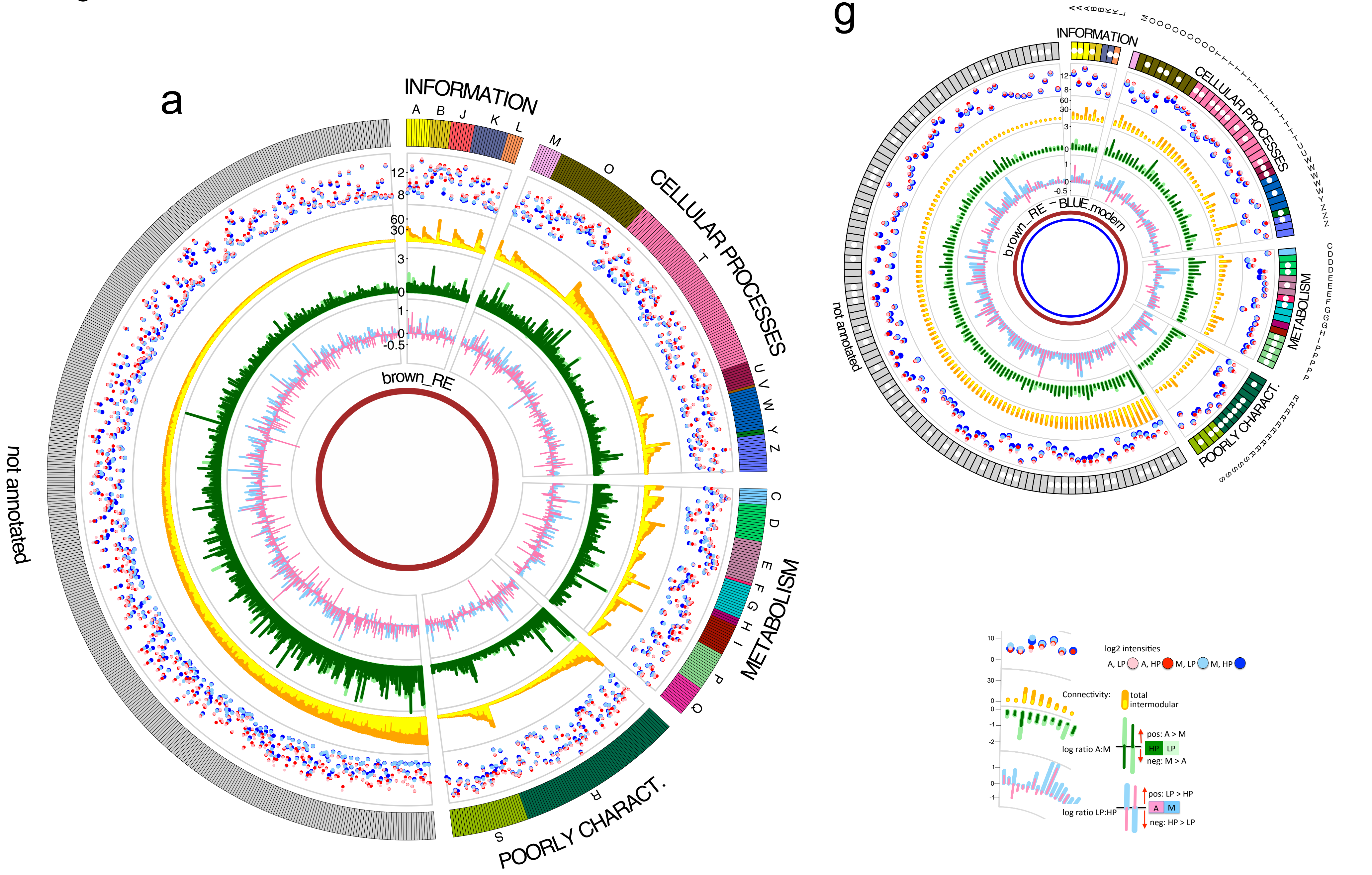
a
INFORMATION
CELLULAR PROCESSES
METABOLISM
POORLY CHARACT.
not annotated
brown_RE
g
brown_RE – BLUE.modern
log2 intensities
A, LP
A, HP
M, LP
M, HP
Connectivity:
total
intermodular
log ratio A:M
pos: A > M
HP
LP
neg: M > A
log ratio LP:HP
pos: LP > HP
A
M
neg: HP > LP

### Figure S5

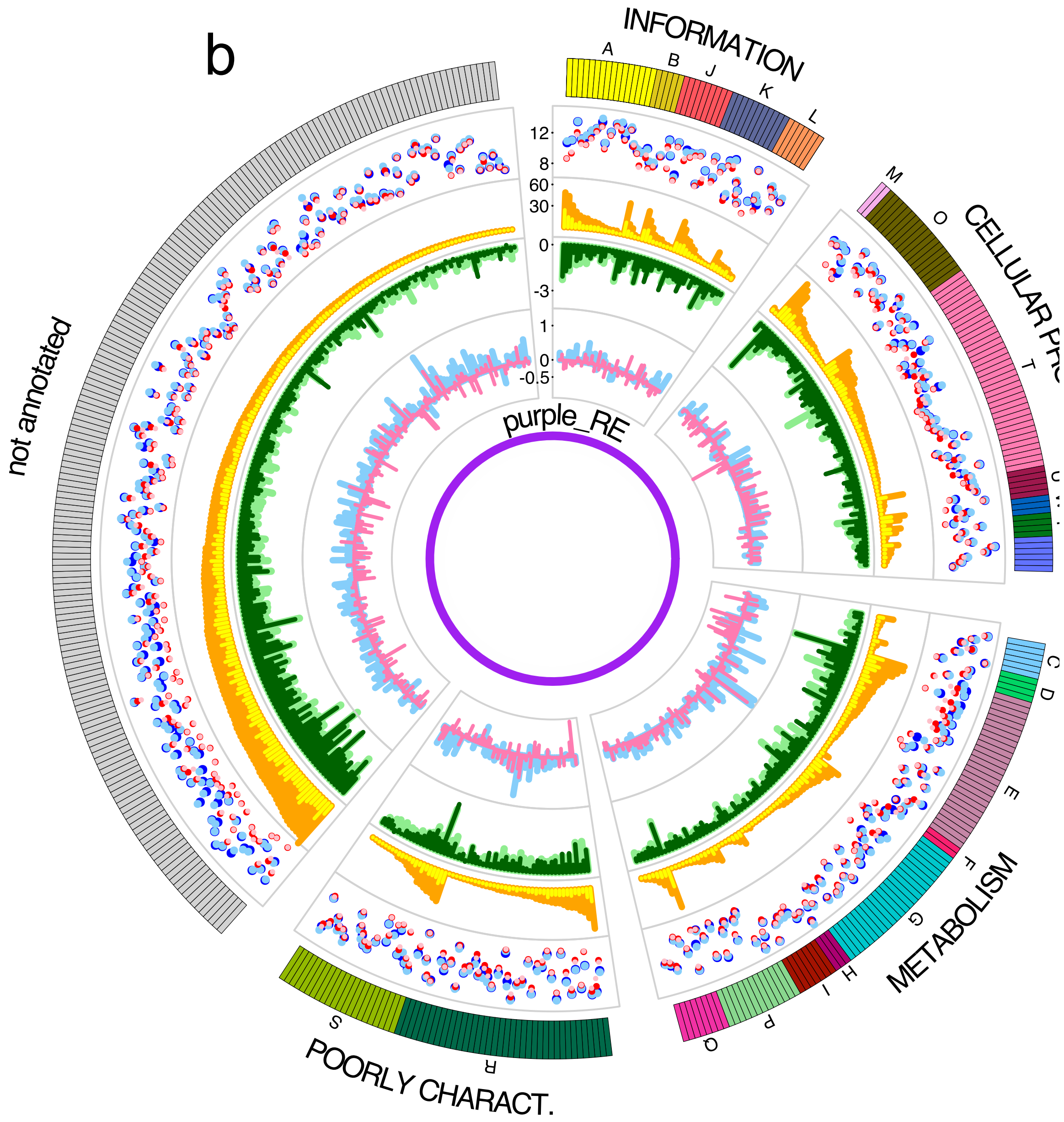
b
INFORMATION
A
B
J
K
L
CELLULAR PR
M
O
T
U
not annotated
purple_RE
METABOLISM
C
D
E
F
G
H
I
P
Q
POORLY CHARACT.
S
R
12
8
60
30
0
-3
1
0
-0.5

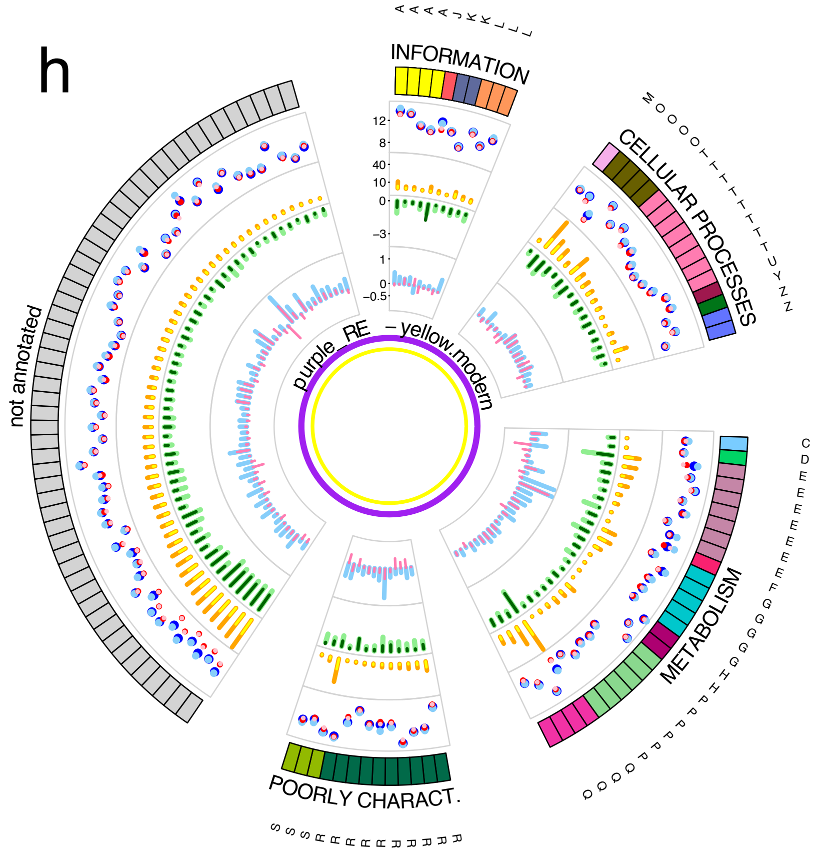
h
INFORMATION
CELLULAR PROCESSES
METABOLISM
POORLY CHARACT.
not annotated
purple_RE – yellow_modern
12
8
40
10
0
-3
1
0
-0.5

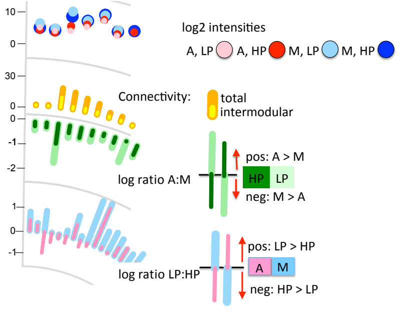
log2 intensities
A, LP
A, HP
M, LP
M, HP
Connectivity:
total
intermodular
log ratio A:M
pos: A > M
HP
LP
neg: M > A
log ratio LP:HP
pos: LP > HP
A
M
neg: HP > LP
10
0
30
0
0
-1
-2
1
0
-1

Figure S5

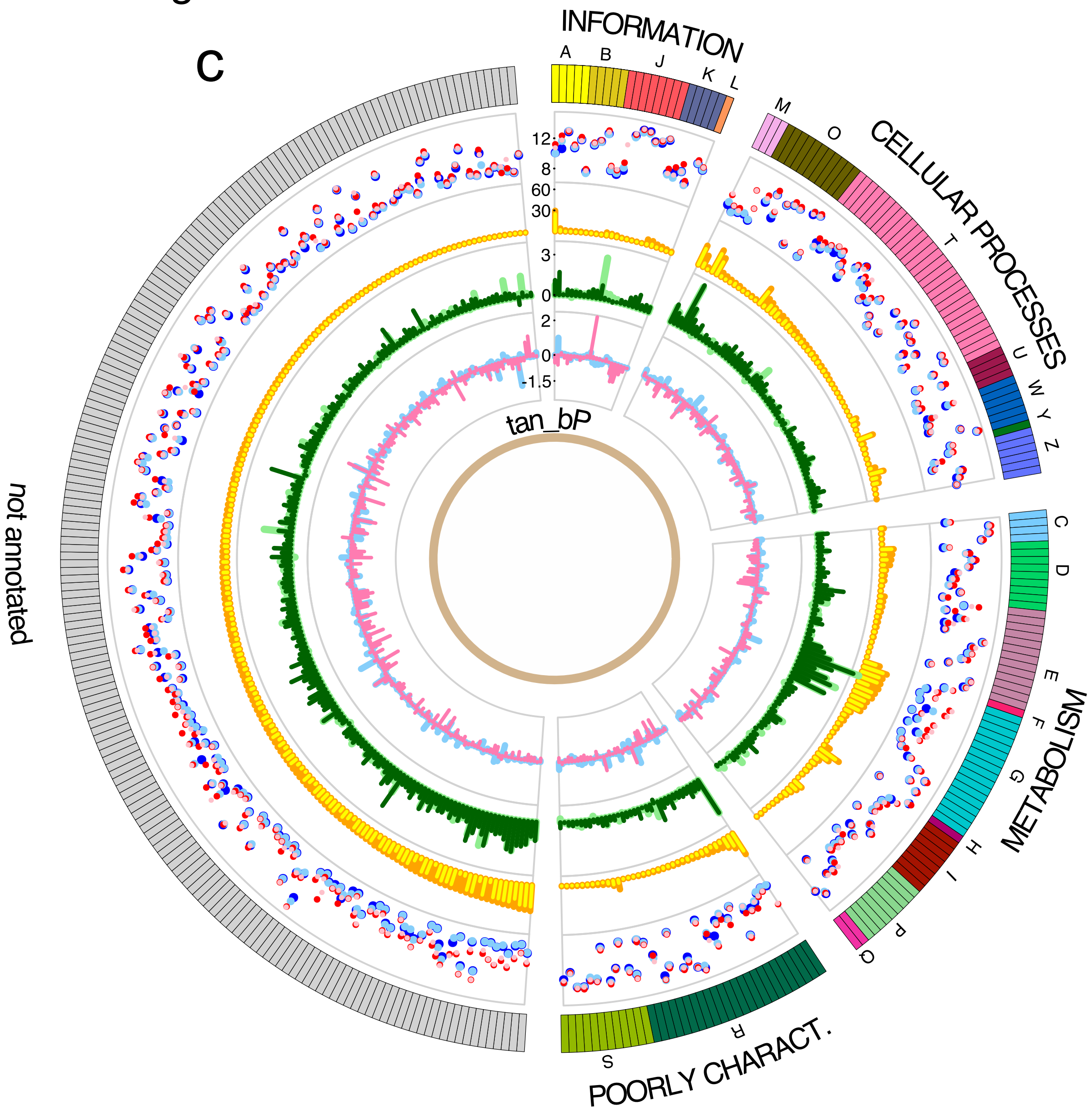

c
INFORMATION
A
B
J
K
L
CELLULAR PROCESSES
M
O
T
U
W
Y
Z
METABOLISM
C
D
E
F
G
H
I
P
Q
POORLY CHARACT.
R
S
not annotated
tan_bP
12
8
60
30
3
0
2
0
-1.5


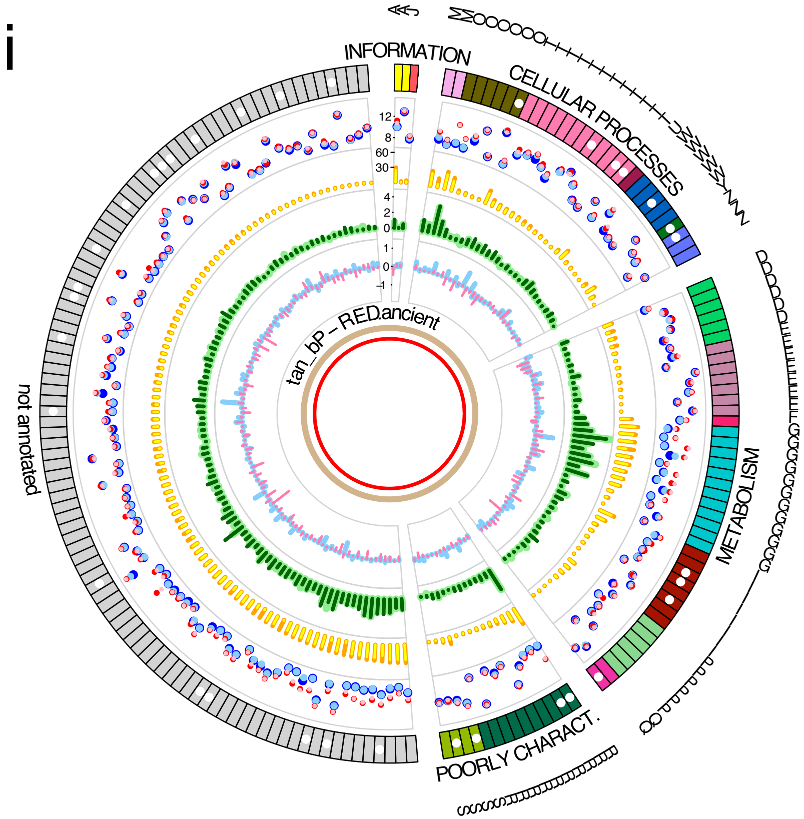

i
INFORMATION
CELLULAR PROCESSES
METABOLISM
POORLY CHARACT.
not annotated
tan_bP - RED.ancient


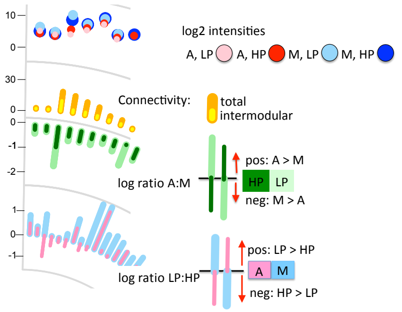

log2 intensities
A, LP
A, HP
M, LP
M, HP
Connectivity:
total
intermodular
log ratio A:M
pos: A > M
HP
LP
neg: M > A
log ratio LP:HP
pos: LP > HP
A
M
neg: HP > LP

Figure S5

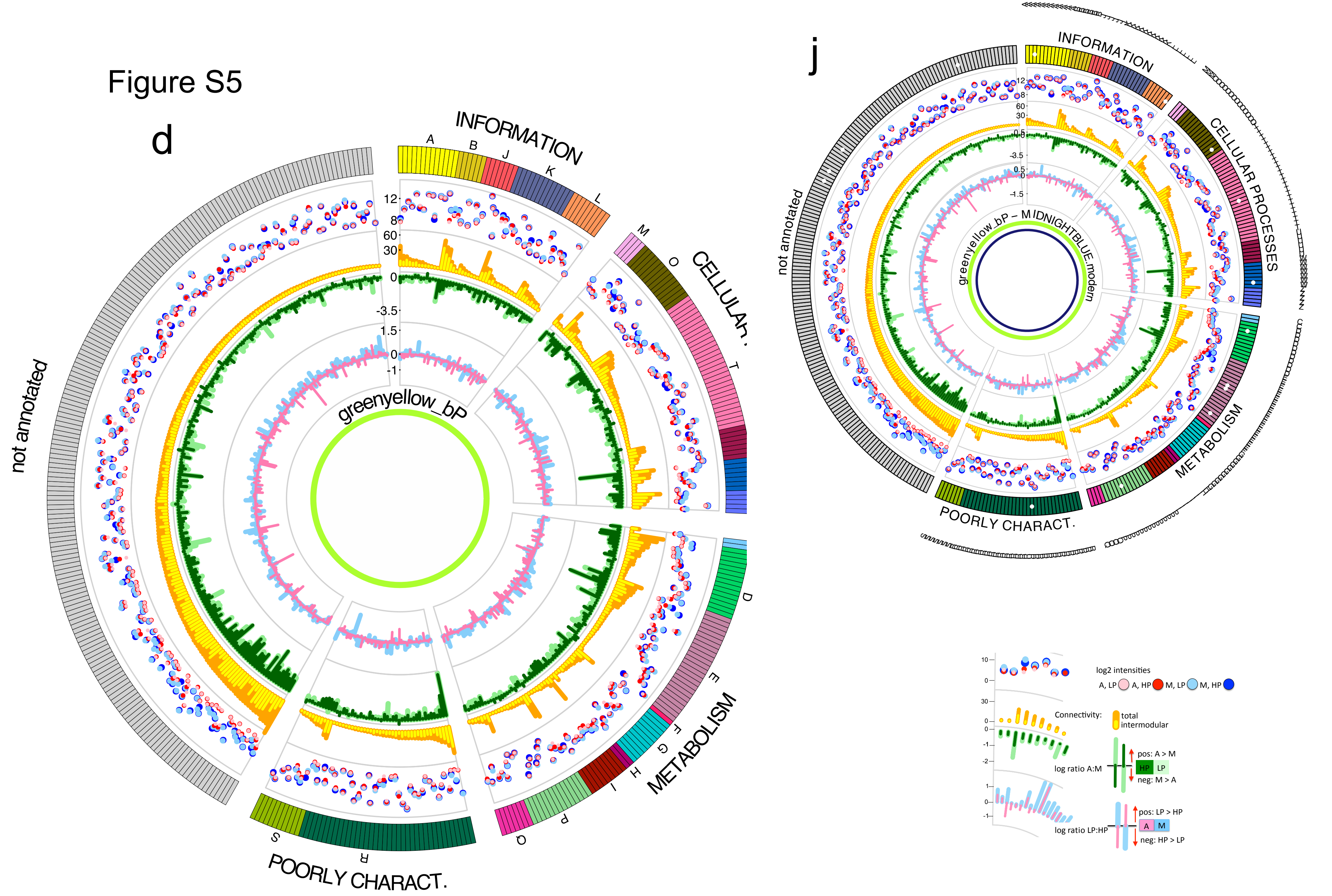

d
INFORMATION
A
B
J
K
L
M
O
CELLULAR
T
D
E
F
G
H
I
METABOLISM
P
Q
S
R
POORLY CHARACT.
not annotated
greenyellow_bP
12
8
60
30
0
-3.5
1.5
0
-1
j
INFORMATION
CELLULAR PROCESSES
METABOLISM
POORLY CHARACT.
not annotated
greenyellow_bP – MIDNIGHTBLUE_modern
log2 intensities
A, LP
A, HP
M, LP
M, HP
Connectivity:
total
intermodular
log ratio A:M
pos: A > M
HP
LP
neg: M > A
log ratio LP:HP
pos: LP > HP
A
M
neg: HP > LP

### Figure S5

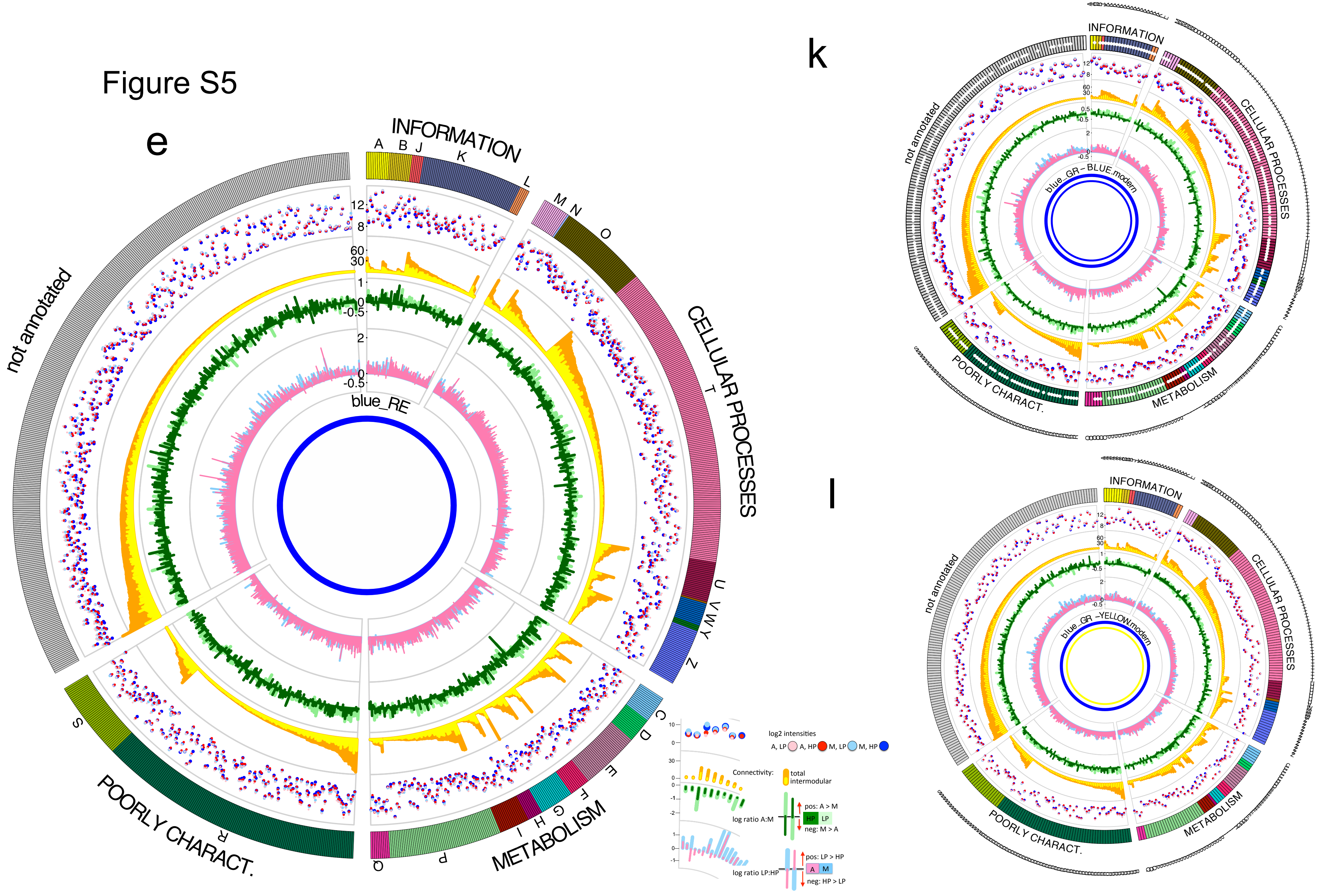

e
INFORMATION
A
B
J
K
L
M
N
O
CELLULAR PROCESSES
T
U
V
W
Y
Z
C
D
E
F
G
H
I
METABOLISM
P
Q
R
S
POORLY CHARACT.
not annotated
blue_RE
12
8
60
30
1
0
-0.5
2
10
log2 intensities
A, LP
A, HP
M, LP
M, HP
Connectivity:
total
intermodular
log ratio A:M
pos: A > M
HP
LP
neg: M > A
log ratio LP:HP
pos: LP > HP
A
M
neg: HP > LP
k
INFORMATION
CELLULAR PROCESSES
METABOLISM
POORLY CHARACT.
not annotated
blue_GR – BLUE.modern
l
INFORMATION
CELLULAR PROCESSES
METABOLISM
POORLY CHARACT.
not annotated
blue_GR –YELLOW.modern

### Figure S5

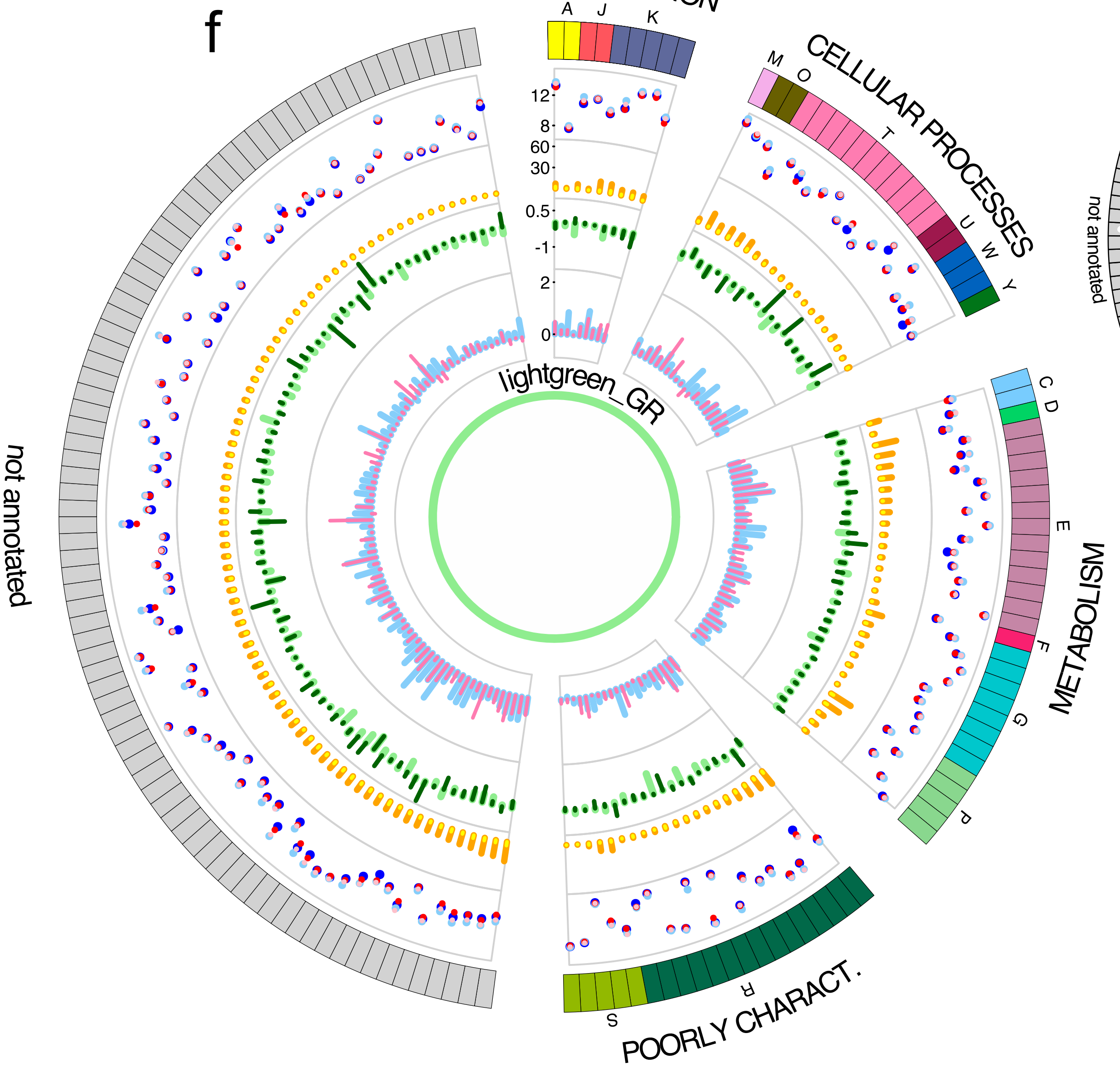
f
INFORMATION
A
J
K
CELLULAR PROCESSES
M
O
T
U
W
Y
METABOLISM
C
D
E
F
G
P
POORLY CHARACT.
S
R
not annotated
lightgreen_GR
12
8
60
30
0.5
-1
2
0

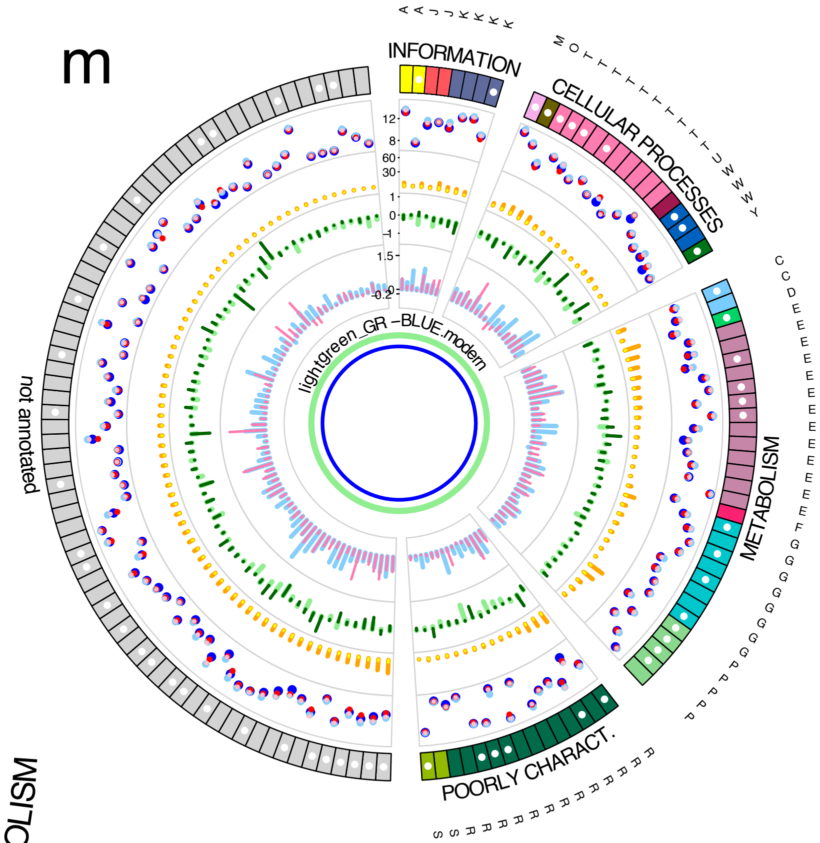
m
INFORMATION
CELLULAR PROCESSES
METABOLISM
POORLY CHARACT.
not annotated
lightgreen_GR –BLUE.modern

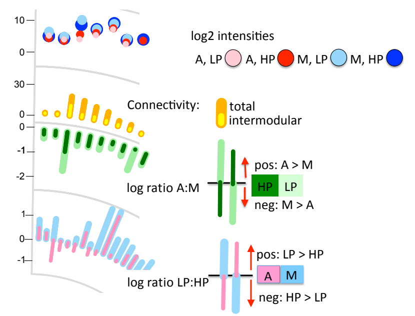
log2 intensities
A, LP
A, HP
M, LP
M, HP
Connectivity:
total
intermodular
log ratio A:M
pos: A > M
HP
LP
neg: M > A
log ratio LP:HP
pos: LP > HP
A
M
neg: HP > LP

Figure S6

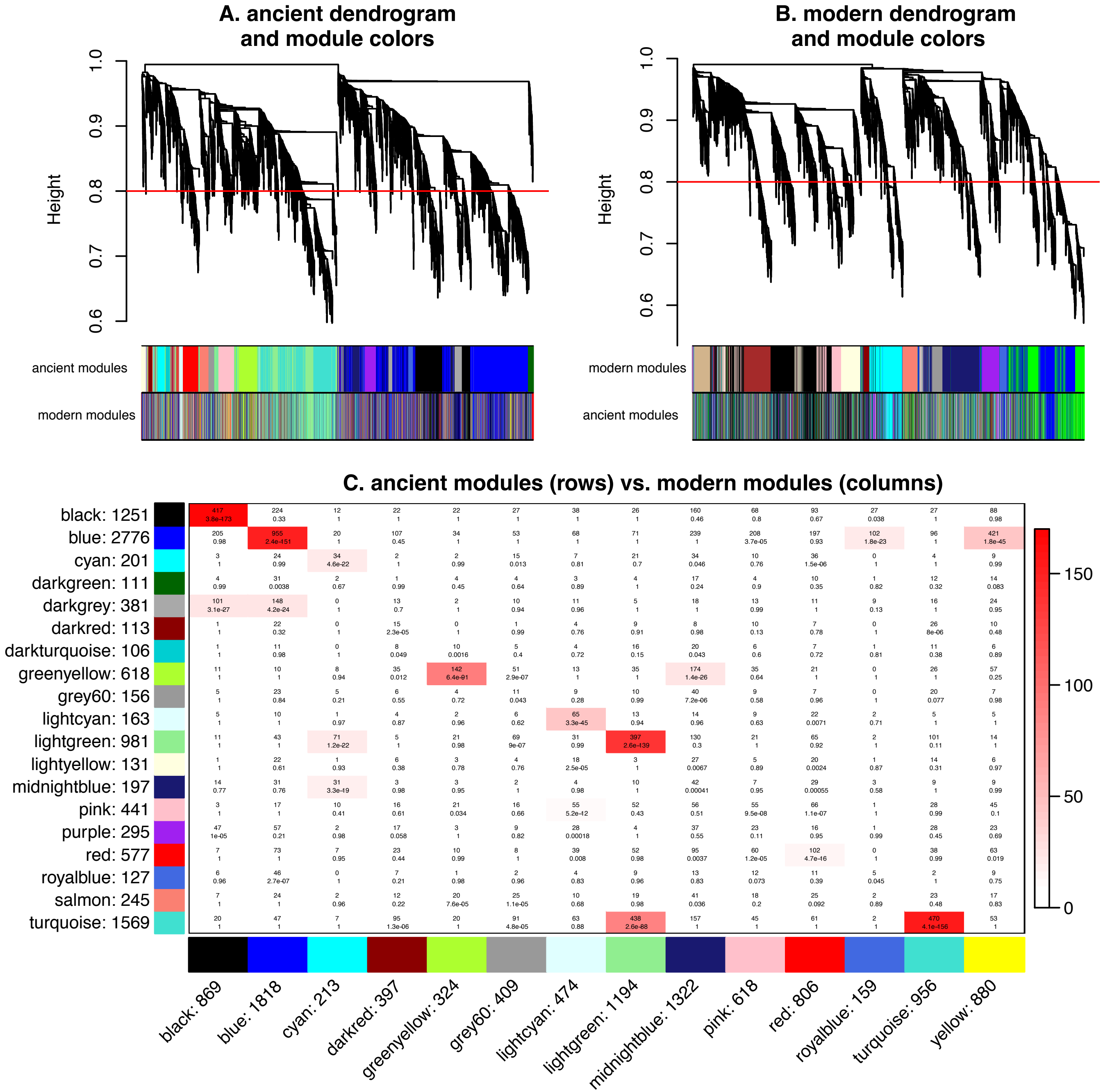
A. ancient dendrogram and module colors
B. modern dendrogram and module colors
Height
0.6
0.7
0.8
0.9
1.0
ancient modules
modern modules
C. ancient modules (rows) vs. modern modules (columns)
black: 1251
blue: 2776
cyan: 201
darkgreen: 111
darkgrey: 381
darkred: 113
darkturquoise: 106
greenyellow: 618
grey60: 156
lightcyan: 163
lightgreen: 981
lightyellow: 131
midnightblue: 197
pink: 441
purple: 295
red: 577
royalblue: 127
salmon: 245
turquoise: 1569
black: 869
blue: 1818
cyan: 213
darkred: 397
greenyellow: 324
grey60: 409
lightcyan: 474
lightgreen: 1194
midnightblue: 1322
pink: 618
red: 806
royalblue: 159
turquoise: 956
yellow: 880
0
50
100
150

### Figure S7

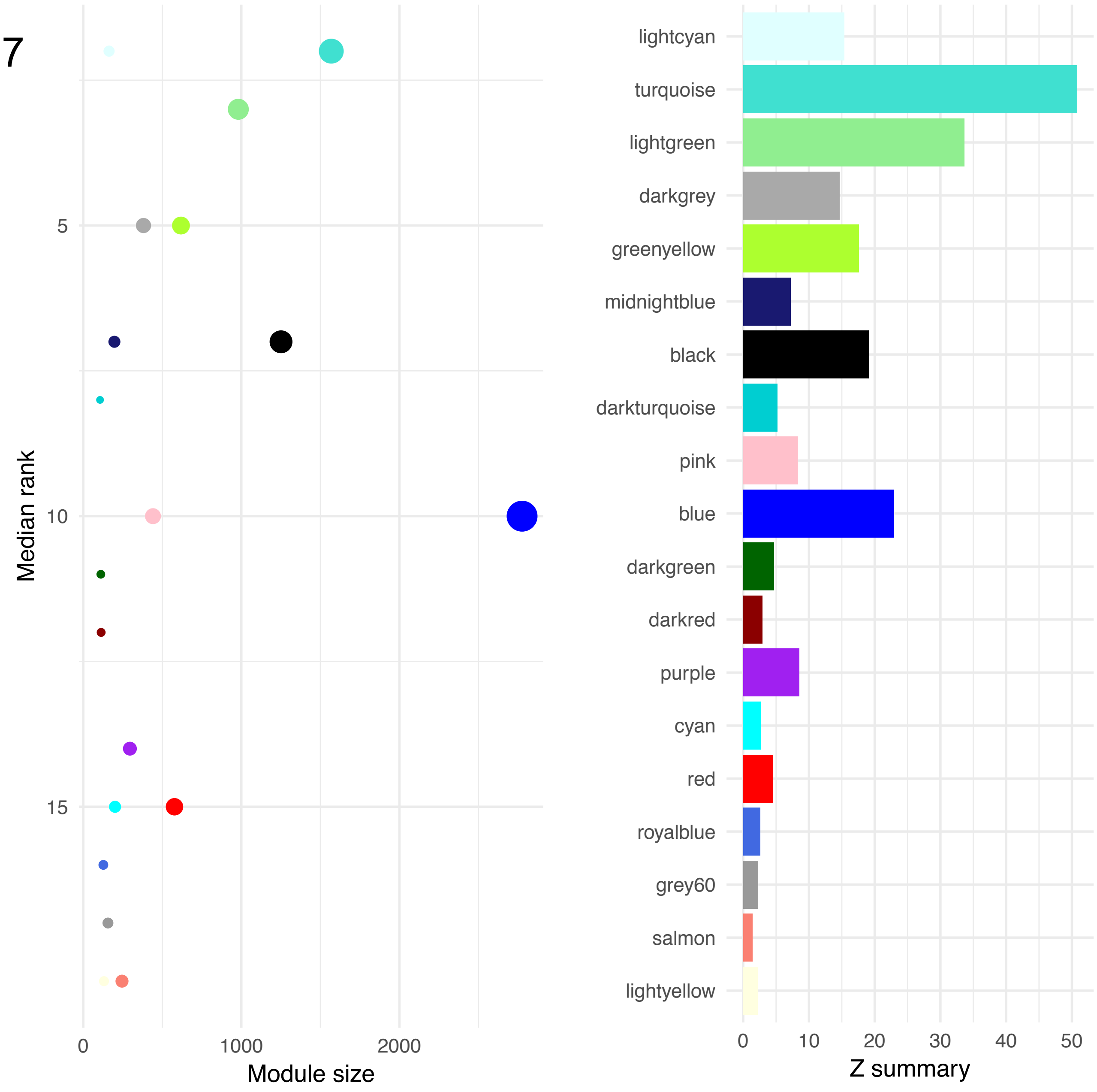

Median rank
5
10
15
0
1000
2000
Module size
lightcyan
turquoise
lightgreen
darkgrey
greenyellow
midnightblue
black
darkturquoise
pink
blue
darkgreen
darkred
purple
cyan
red
royalblue
grey60
salmon
lightyellow
0
10
20
30
40
50
Z summary

### Figure S8

#### Enrichment of KOG groups in the modules that were **preserved** between ancient and modern clone networks

| preserved module | turquoise | | lightgreen | | greenyellow | | lightcyan | |
|---|---|---|---|---|---|---|---|---|
| ancient (A) / modern (M) | A | M | A | M | A | M | A | M |
| number of genes | 1569 | 956 | 981 | 1194 | 618 | 324 | 163 | 474 |
| **Information storage and processing** | 1.43 | | 1.84 | 1.42 | 1.71 | 1.67 | | |
| [A] RNA processing & modification | | | | | 2.25 | 2.32 | | |
| [B] Chromatin structure & dynamics | | | 1.89 | | | | | |
| [J] Translation, ribosomal structure & biogenesis | | 0.42 | 2.67 | | 4.99 | 2.76 | | |
| [K] Transcription | 1.40 | | | 1.62 | 0.49 | | | |
| [L] Replication, recombination & repair | 1.51 | | 1.80 | | | | | 0.10 |
| **Cellular processes and signalling** | 0.72 | 0.76 | 0.73 | 0.70 | 0.81 | | | |
| [M] Cell wall/membrane/envelope biogenesis | | | | | | | | 2.90 |
| [N] Cell motility | | | | | | | | |
| [O] Posttranslational modification, protein turnover, chaperones | | | | | 2.51 | 2.57 | | |
| [T] Signal transduction mechanisms | 0.64 | | 0.70 | 0.68 | 0.19 | 0.29 | | |
| [U] Intracellular trafficking, secretion and vesicular transport | | | | | 2.24 | | | |
| [V] Defense mechanisms | | | | | 0.16 | | | 2.16 |
| [W] Extracellular structures | 0.39 | 0.35 | | | 0.13 | | | |
| [Y] Nuclear structure | | | | | | | | |
| {Z] Cytoskeleton | 0.54 | | | | | | | |
| **Metabolism** | | | | | 0.70 | | 1.81 | 1.39 |
| [C] Energy production and conversion | | | | 0.26 | | | | |
| [D] Cell cycle control, cell division, chromosome partitioning | 3.04 | 3.23 | 2.52 | 3.16 | | | | |
| [E] Amino acid transport and metabolism | 0.60 | | 0.50 | | 0.45 | | 4.18 | |
| [F] Nucleotide transport and metabolism | | | | | 2.37 | | | |
| [G] Carbohydrate transport and metabolism | 0.50 | | 0.46 | | | | | |
| [I] Lipid transport and metabolism | | | | | 0.48 | | | 3.02 |
| [P] Inorganic ion transport and metabolism | | | 0.30 | | 0.38 | 0.19 | | |
| [Q] Second. metabolites biosynth., transport & catabolism | | | | | | | | |
| **Poorly characterized** | | | | | | | 0.51 | 0.50 |
| [R] General function prediction only | | | | | 0.73 | 0.58 | | 0.59 |
| [S] Function unknown | | | | | 2.01 | 1.69 | | 0.37 |

#### Enrichment in modules that were **not preserved** between ancient and modern clone networks

| not preserved modules | salmon | | grey60 | | lightyellow | | royalblue | |
|---|---|---|---|---|---|---|---|---|
| ancient (A) / modern (M) | A | M | A | M | A | M | A | M |
| number of genes | 245 | N/A | 156 | 409 | 131 | N/A | 127 | 159 |
| **Information storage and processing** | 1.67 | | | 1.96 | | | | |
| [A] RNA processing & modification | | | | | | | | |
| [B] Chromatin structure & dynamics | | | | | | | | |
| [J] Translation, ribosomal structure & biogenesis | 3.11 | | | 8.36 | | | | |
| [K] Transcription | | | | | | | | |
| [L] Replication, recombination & repair | | | | | | | | |
| **Cellular processes and signalling** | | | | 0.73 | | | | |
| [M] Cell wall/membrane/envelope biogenesis | | | | | | | | |
| [N] Cell motility | | | | | | | | |
| [O] Posttranslational modification, protein turnover, chaperones | | | | 1.86 | | | | |
| [T] Signal transduction mechanisms | | | | 0.59 | | | | |
| [U] Intracellular trafficking, secretion and vesicular transport | | | | | | | | |
| [V] Defense mechanisms | | | | | | | | |
| [W] Extracellular structures | | | | | | | | |
| [Y] Nuclear structure | | | | | | | | |
| {Z] Cytoskeleton | | | | | | | | |
| **Metabolism** | | | | | | | | |
| [C] Energy production and conversion | | | | 3.38 | | | | |
| [D] Cell cycle control, cell division, chromosome partitioning | | | | | | | | |
| [E] Amino acid transport and metabolism | | | | | | | | |
| [F] Nucleotide transport and metabolism | | | | | | | | |
| [G] Carbohydrate transport and metabolism | | | | | | | | |
| [I] Lipid transport and metabolism | | | | | | | | |
| [P] Inorganic ion transport and metabolism | | | | | | | | |
| [Q] Second. metabolites biosynth., transport & catabolism | | | | | | | | |
| **Poorly characterized** | | | | | | | | |
| [R] General function prediction only | | | | 0.55 | | | | |
| [S] Function unknown | 2.28 | | | 1.95 | | | | |

Figure S9

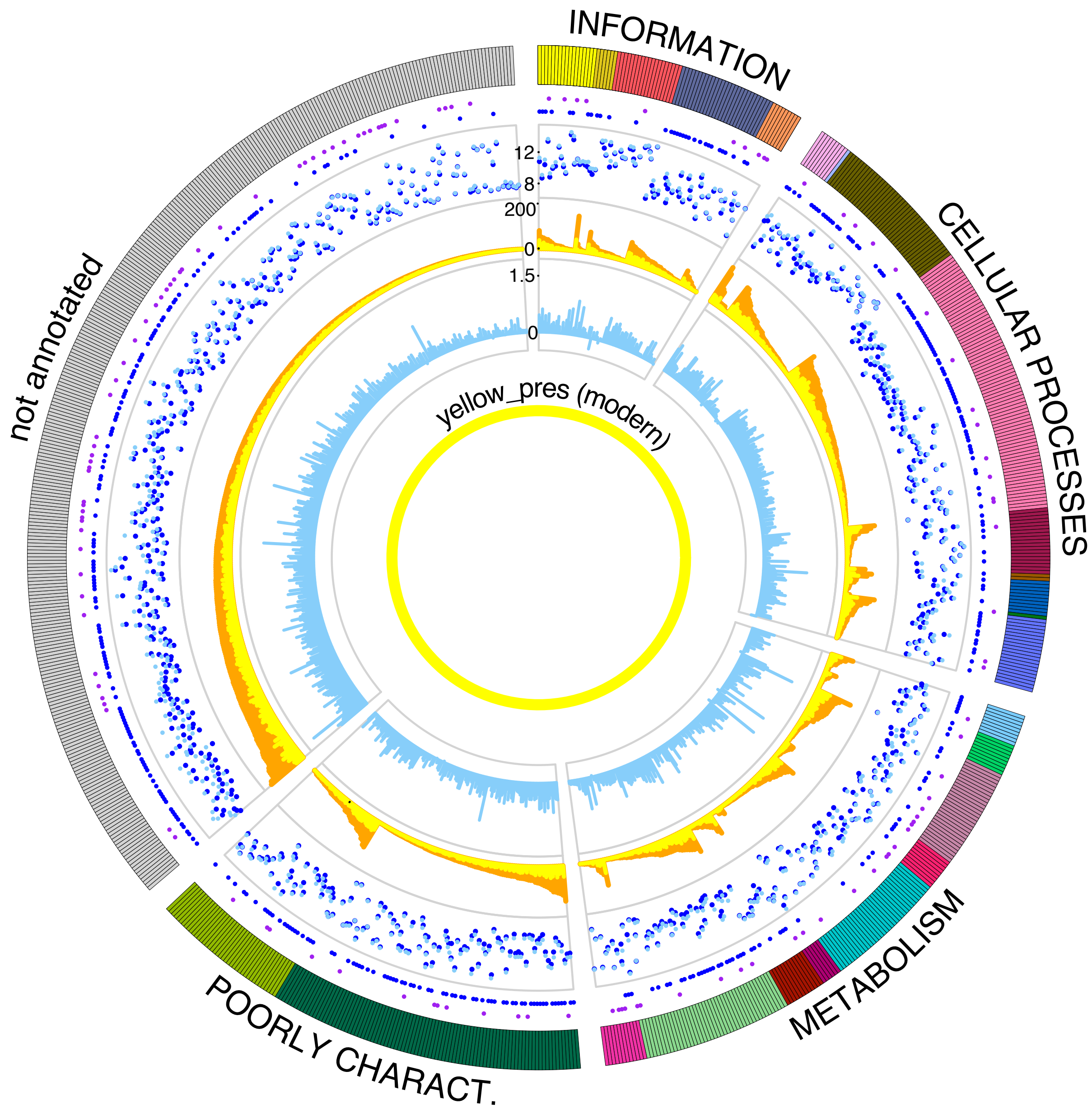
INFORMATION
CELLULAR PROCESSES
METABOLISM
POORLY CHARACT.
not annotated
yellow_pres (modern)
12
8
200
0
1.5
0

### Figure S10

#### Percentage of RE genes in preservation network modules

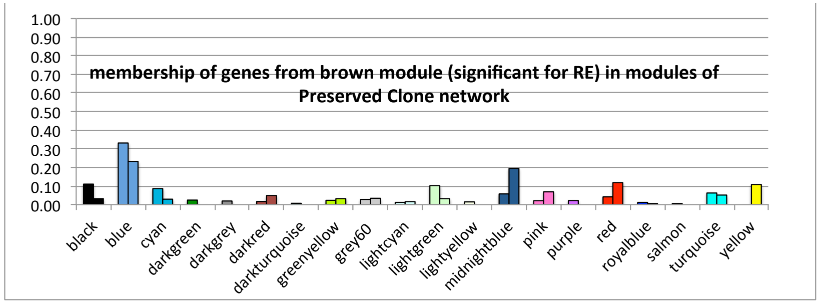


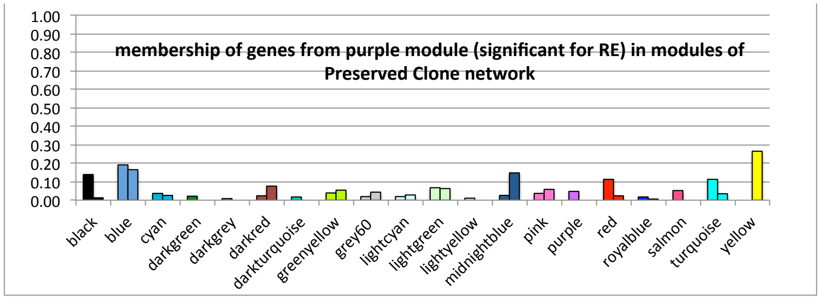


#### Percentage of bP genes in preservation network modules

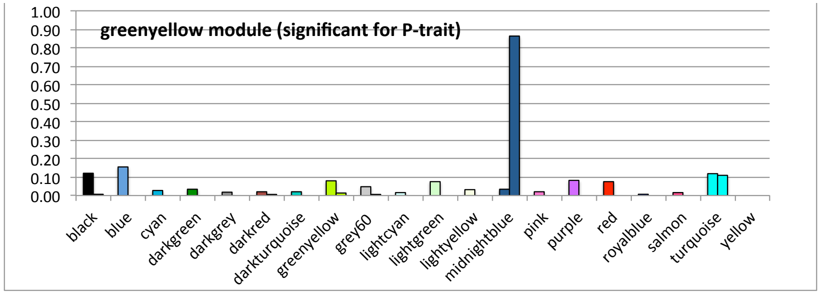


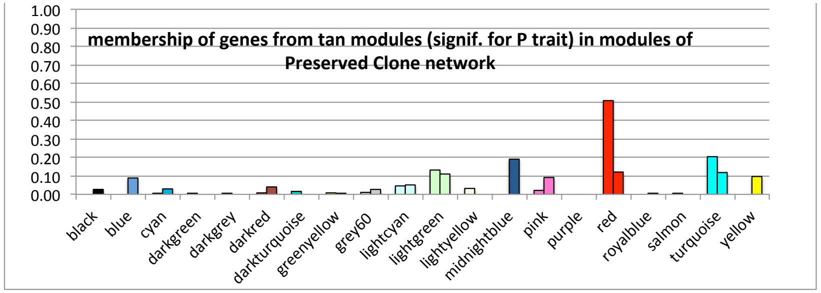


#### Percentage of GR genes in preservation network modules

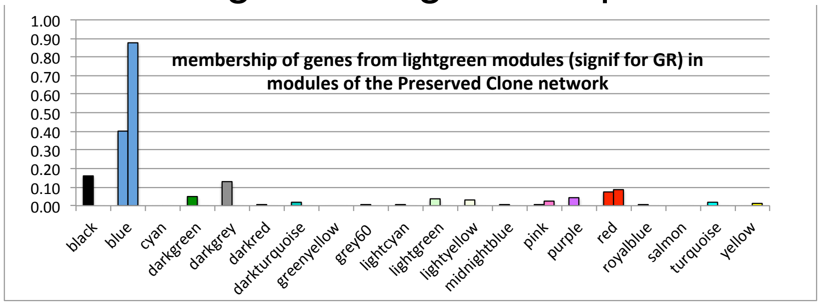


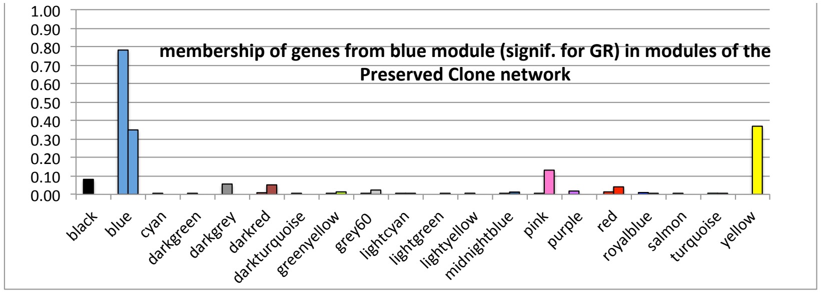

### Table S1

| Module | Size | Density | Centralization | Heterogeneity | Connectivity | Scaled.connect. | Clustering.coeff. | Max.adj.ratio |
|---|---|---|---|---|---|---|---|---|
| black | 920 | 0.078 | 0.129 | 0.633 | 71.793 | 0.379 | 0.153 | 0.16 |
| blue | 1106 | 0.066 | 0.109 | 0.541 | 72.766 | 0.377 | 0.112 | 0.125 |
| brown | 815 | 0.075 | 0.129 | 0.535 | 60.709 | 0.367 | 0.13 | 0.146 |
| cyan | 317 | 0.134 | 0.149 | 0.555 | 42.292 | 0.475 | 0.224 | 0.226 |
| green | 702 | 0.056 | 0.067 | 0.407 | 39.009 | 0.454 | 0.086 | 0.113 |
| greenyellow | 439 | 0.102 | 0.15 | 0.546 | 44.53 | 0.406 | 0.177 | 0.185 |
| grey60 | 165 | 0.156 | 0.145 | 0.437 | 25.542 | 0.52 | 0.229 | 0.24 |
| lightcyan | 219 | 0.101 | 0.105 | 0.413 | 22.068 | 0.493 | 0.15 | 0.172 |
| lightgreen | 162 | 0.084 | 0.085 | 0.384 | 13.531 | 0.501 | 0.123 | 0.152 |
| magenta | 576 | 0.095 | 0.151 | 0.657 | 54.823 | 0.389 | 0.181 | 0.184 |
| pink | 686 | 0.076 | 0.132 | 0.719 | 52.396 | 0.368 | 0.165 | 0.165 |
| purple | 461 | 0.082 | 0.132 | 0.542 | 37.703 | 0.385 | 0.142 | 0.152 |
| red | 695 | 0.092 | 0.126 | 0.46 | 63.809 | 0.423 | 0.14 | 0.155 |
| salmon | 322 | 0.049 | 0.051 | 0.368 | 15.765 | 0.49 | 0.085 | 0.119 |
| tan | 373 | 0.066 | 0.133 | 0.728 | 24.618 | 0.334 | 0.154 | 0.159 |
| turquoise | 1691 | 0.085 | 0.146 | 0.687 | 144.169 | 0.37 | 0.175 | 0.169 |
| yellow | 790 | 0.075 | 0.137 | 0.622 | 59.375 | 0.355 | 0.15 | 0.157 |

### Table S2

| Ancient Module | Size | Density | Centralization | Heterogeneity | Connectivity | Scaled.connect. | Clustering.coeff. | Max.adj.ratio |
|---|---|---|---|---|---|---|---|---|
| black | 1251 | 0.12 | 0.16 | 0.53 | 147.73 | 0.43 | 0.20 | 0.22 |
| blue | 2776 | 0.08 | 0.13 | 0.64 | 220.63 | 0.38 | 0.16 | 0.18 |
| cyan | 201 | 0.09 | 0.12 | 0.48 | 18.15 | 0.44 | 0.15 | 0.17 |
| darkgreen | 111 | 0.09 | 0.09 | 0.45 | 10.17 | 0.52 | 0.16 | 0.20 |
| darkgrey | 381 | 0.17 | 0.18 | 0.50 | 62.96 | 0.48 | 0.26 | 0.27 |
| darkred | 113 | 0.09 | 0.11 | 0.47 | 9.84 | 0.45 | 0.14 | 0.17 |
| darkturquoise | 106 | 0.11 | 0.14 | 0.41 | 11.49 | 0.45 | 0.16 | 0.19 |
| greenyellow | 618 | 0.12 | 0.12 | 0.41 | 71.32 | 0.49 | 0.17 | 0.20 |
| grey60 | 156 | 0.16 | 0.16 | 0.39 | 25.22 | 0.50 | 0.22 | 0.24 |
| lightcyan | 163 | 0.16 | 0.15 | 0.50 | 25.06 | 0.51 | 0.24 | 0.25 |
| lightgreen | 981 | 0.10 | 0.15 | 0.63 | 96.58 | 0.39 | 0.19 | 0.19 |
| lightyellow | 131 | 0.08 | 0.11 | 0.52 | 10.28 | 0.42 | 0.16 | 0.20 |
| midnightblue | 197 | 0.10 | 0.11 | 0.41 | 19.45 | 0.47 | 0.15 | 0.18 |
| pink | 441 | 0.14 | 0.14 | 0.44 | 63.58 | 0.50 | 0.22 | 0.23 |
| purple | 295 | 0.13 | 0.14 | 0.40 | 39.46 | 0.50 | 0.20 | 0.23 |
| red | 577 | 0.13 | 0.17 | 0.55 | 74.58 | 0.43 | 0.22 | 0.23 |
| royalblue | 127 | 0.20 | 0.18 | 0.36 | 24.76 | 0.52 | 0.26 | 0.28 |
| salmon | 245 | 0.15 | 0.13 | 0.39 | 37.01 | 0.53 | 0.21 | 0.24 |
| turquoise | 1569 | 0.11 | 0.16 | 0.63 | 172.74 | 0.41 | 0.21 | 0.21 |

| Modern Module | Size | Density | Centralization | Heterogeneity | Connectivity | Scaled.connect. | Clustering.coeff. | Max.adj.ratio |
|---|---|---|---|---|---|---|---|---|
| black | 869 | 0.11 | 0.16 | 0.59 | 95.84 | 0.41 | 0.21 | 0.22 |
| blue | 1818 | 0.07 | 0.12 | 0.58 | 127.49 | 0.37 | 0.14 | 0.16 |
| cyan | 213 | 0.10 | 0.11 | 0.39 | 21.17 | 0.47 | 0.14 | 0.18 |
| darkred | 397 | 0.07 | 0.09 | 0.44 | 27.99 | 0.44 | 0.12 | 0.15 |
| greenyellow | 324 | 0.11 | 0.10 | 0.36 | 33.89 | 0.53 | 0.15 | 0.18 |
| grey60 | 409 | 0.08 | 0.09 | 0.50 | 33.30 | 0.47 | 0.14 | 0.17 |
| lightcyan | 474 | 0.11 | 0.15 | 0.56 | 51.70 | 0.42 | 0.20 | 0.22 |
| lightgreen | 1194 | 0.12 | 0.16 | 0.61 | 138.10 | 0.42 | 0.22 | 0.22 |
| midnightblue | 1322 | 0.08 | 0.14 | 0.68 | 107.09 | 0.37 | 0.17 | 0.18 |
| pink | 618 | 0.11 | 0.16 | 0.66 | 66.56 | 0.40 | 0.21 | 0.20 |
| red | 806 | 0.13 | 0.18 | 0.56 | 106.76 | 0.43 | 0.23 | 0.23 |
| royalblue | 159 | 0.10 | 0.11 | 0.39 | 15.19 | 0.47 | 0.14 | 0.17 |
| turquoise | 956 | 0.12 | 0.17 | 0.61 | 117.12 | 0.42 | 0.23 | 0.23 |
| yellow | 880 | 0.10 | 0.14 | 0.56 | 84.35 | 0.40 | 0.17 | 0.18 |
